# Microbial dark matter driven degradation of carbon fiber polymer composites

**DOI:** 10.1101/2020.04.05.024463

**Authors:** Adam M. Breister, Muhammad A. Imam, Zhichao Zhou, Karthik Anantharaman, Pavana Prabhakar

**Author notes:** Equal Contribution (Adam M. Breister, Muhammad A. Imam).

## Abstract

Polymer composites have become attractive for structural applications in the built environment due to their lightweight and high strength properties but can suffer from degradation due to environmental factors. While impacts of abiotic factors like temperature and moisture are well studied, little is known about the influence of naturally occurring microbial communities on their structural integrity. Here we apply complementary time-series multi-omics of biofilms growing on polymer composites and materials characterization to elucidate, for the first time, the processes driving their degradation. We measured a reduction in mechanical properties due to molecular chain breakage and reconstructed 121 microbial genomes to describe microbial diversity and pathways associated with their degradation. The composite microbiome is dominated by four bacterial groups including the Candidate Phyla Radiation that possess pathways for breakdown of acrylate, esters, and bisphenol, abundant in composites. Overall, we provide a foundation for understanding interactions of next-generation structural materials with their natural environment that can predict their durability and drive future designs.

## Introduction

Fiber-reinforced polymer composite materials (referred to as polymer composites) have become an attractive option for infrastructure due to their lightweight, high stiffness and high strength to weight ratios^1^, which are favorable properties for enabling high fuel efficiency and towards increasing mobility. In addition, polymer composites have extremely high corrosion resistance compared to conventional materials like steel which are easily corroded. Therefore, although initial installed costs of polymer composites and steel are comparable, life-cycle costs of polymer composites are expected to be significantly lower. Currently, polymer composites have a prevalent use in a large range of infrastructure^2^ including pipelines, utility poles, prefabricated pavements, renewable energy harvesting, chimneys or flues, rapidly deployable housing, decking for marine and naval structures and advanced retrofitting. Recent studies^3,4^ have demonstrated that polymer composites, when designed appropriately, can provide a cost-effective alternative to current structural materials used for primary loading-bearing elements in civil infrastructure. Specific civil infrastructural applications of polymer composites include but are not limited to retrofitting of existing concrete structures, strengthening of steel structures, as internal reinforcing rods in reinforced concrete structures, in new construction as structural members^5^, in supporting frame structures^6^, multi-storey office^7^ and residential buildings^8^, bridge decks for pedestrian and highway bridge superstructures^9^, electricity transmission towers^10^, and composite piles used for foundation construction^11^. Given their prevalent use in current and future human establishments, it is of utmost importance to understand and elucidate their long-term durability and survivability in natural environments.

Polymer composites are composed of several individual constituent materials with unique properties that are combined together to achieve improved physical properties as compared to the individual materials. They typically consist of two distinct components, a matrix and reinforcing materials. Reinforcing materials such as fibers provide strength to the composite, while the matrix acts as a binding agent to hold the composite together. A common type of polymer composite are fiber-reinforced polymer composites where fibers such as those made of carbon act as reinforcements in a polymer matrix, and polymers such as epoxy, vinyl ester and polyesters are used as matrix. Carbon fibers are widely used for reinforcement due to their high-strength and lightweight properties, and vinyl ester is widely used as binding matrices due to better resistance to moisture absorption and UV radiation compared to polyesters and other binding agents^12,13^. Nevertheless, polymer composites are susceptible to degradation by the chemical, physical and biological stressors in the environment. Key environmental factors that can influence the durability of polymer composites include moisture, temperature, pH, salinity, sustained stresses, and microbes. The response of polymer composites to a few of these environmental factors, like temperature, moisture, pH, freeze-thaw cycles, acting independently or in combination have been well studied^14–19^. However, the influence of microbial interactions on the durability and longevity of polymer composites has seldom been studied. In this current study, we investigate the durability and survivability of a commonly used polymer composite with woven carbon fibers as the reinforcement in vinyl ester polymer matrix.

Previous studies have demonstrated that incubation of binding matrices such as epoxy with individual organisms such as *Pseudomonas spp*. as well as simple consortia of microorganisms can degrade these compounds^20–24^. These microorganisms typically formed a viscoelastic layer or biofilm on the material surface^25–27^. Possible mechanisms for microbial degradation of materials involved the degradation of organic polymers in the binding matrix involving direct attack by acids or enzymes, blistering due to gas evolution, enhanced cracking due to calcareous deposits and gas evolution and polymer destabilization by concentrated chlorides and sulfides^24^.

Among these very limited studies on polymer composites, most have focused only on the influence of single microorganisms on specific compounds such as epoxy in controlled environments. Wagner et al.^24^ exposed polymer composites to four different microbial cultures, namely *Thiobacillus ferroxidans* (a sulfur/iron oxidizing bacterium), *Pseudomonas fluorescens* (a calcareous-depositing bacterium), *Lactococcus lactis* (ammonium and sulfide producing bacterium) and sulfate-reducing bacteria (SRB; producing sulfide from sulfate), on polymer composites. Bacterial colonization of fibers and composites were observed. Hydrogen-producing bacteria appeared to disrupt bonding between fibers and vinyl ester resin in the polymer composites and penetrated the resin at the interface in addition to disrupting fibers and resin due to gas formation within the composite. The strength of polymer composites reduced after exposure to hydrogen sulfide (from *Lactococcus* and SRB) and its corrosive effects. In addition, polymer composites can also be impacted by leaching activities of heterotrophic bacteria that seek carbon from the polymer and production of reactive oxygen species during growth. Carbon is a critical nutrient necessary for microbial growth and organic components in binding matrices such as epoxy can serve as the sole source of carbon for organisms found in soil such as *Rhodococcus rhodochrous* and *Ochrobactrum anthropic*^21^.

In addition to bacteria, a few studies have also investigated the impact of fungi on polymer compsites^23,28^. Investigations using a fungal consortium (*Aspergillus versicolor, Cladosporium cladospoiordes*, and *Chaetomum*) on different polymer composites revealed that all samples were colonized by the consortia with fungal penetration along the fibers accelerating the attack on the binding matrix. It is likely that organic compounds in the polymer composites likely served as carbon and energy sources for the growth of fungi. Though important, these single organism-based studies do not adequately capture the impacts of a complex natural microbial community on polymer composites. Moreover, much of these organisms used in prior studies are not abundant in molecular surveys conducted in nature (for example in soils) and are thus unlikely to be the key drivers of degradation of composites^29^. Therefore, little is known about the diversity and impacts of microbes interacting with polymer composites in nature. Cultivation-independent studies that can comprehensively characterize the microbial community and its deleterious effects on composites are needed to understand their impacts and design mitigation strategies.

Here, we study the impact of a complex natural microbial community on the degradation of polymer composites over time. We performed complementary mechanical and materials characterization and multi-omics analyses to assess the extent and type of degradation of the polymer composites and obtain a genome-informed perspective on the potential role of microorganisms in this process. By utilizing a time-course experiment, we were able to observe a gradual reduction in a number of metrics associated with strength, stiffness, durability, and survivability of polymer composites. Our sophisticated materials characterization approach revealed for the first time, specific mechanisms that underpinned degradation of polymer composites by microorganisms over time. Our microbial community composition and metagenomic analyses support our findings from mechanical and materials characterizations. We observed the presence of a stable biofilm on the polymer composites and utilized genome-resolved analyses to demonstrate the putative roles of poorly studied microbes, including a significant proportion from uncultivated candidate phyla in the degradation of polymer composites. Overall, our study demonstrates that uncultivated microbes in nature possess the ability to degrade widely used polymer composite materials in the built environment.

## Results

### Exposure study of polymer composites: Sampling the polymer composite microbiome and materials characterization

We used microbial community analyses based on 16S ribosomal RNA sequencing and genome-resolved metagenomics, and mechanical and materials characterization based on Thermogravimetric analysis (TGA) and Fourier-transform infrared spectroscopy (FT-IR) to study the impacts of microorganisms on the degradation of polymer composite materials. We fabricated vinyl ester based carbon-fiber reinforced composites in-house using vacuum assisted resin transfer molding process, which is commonly used for fabricating such composites^15,30,31^. These composites were fabricated using layers of woven carbon fiber bundles and stacking them in the thickness direction, followed by infiltration of vinyl ester resin through the dry fabric. Solid composites were obtained upon curing of the resin with the carbon fiber layers. Optical and scanning electron microscopy (SEM) images of the fabricated polymer composite are shown in Figure 1 at different length scales. We inoculated these polymer composites with a soil solution (Soil with deionized water, S + DI) containing soil samples collected from an area adjacent to Lake Mendota, WI, USA (Figure 2). We hypothesized that these soil samples contained a diverse microbial community that would colonize the polymer composites. Two types of control samples were maintained and analyzed over the course of the experiment. First, polymer composites were incubated in autoclaved soil solution with deionized water (S + DI + A). These samples allowed us to measure the impact of soil chemistry (without microorganisms) on the polymer composites. Secondly, polymer composites were incubated in autoclaved DI water (DI+A) to measure the impacts of water on polymer composites (Without microbial and chemical impact). Immediately post-inoculation, we observed the presence of a biofilm on the polymer composites incubated in soil with deionized water. No biofilms were observed in the two control samples. The experimental and control groups were observed over a time course of 24 weeks and samples for microbial and materials characterization were collected every two weeks.

**Figure 1.**
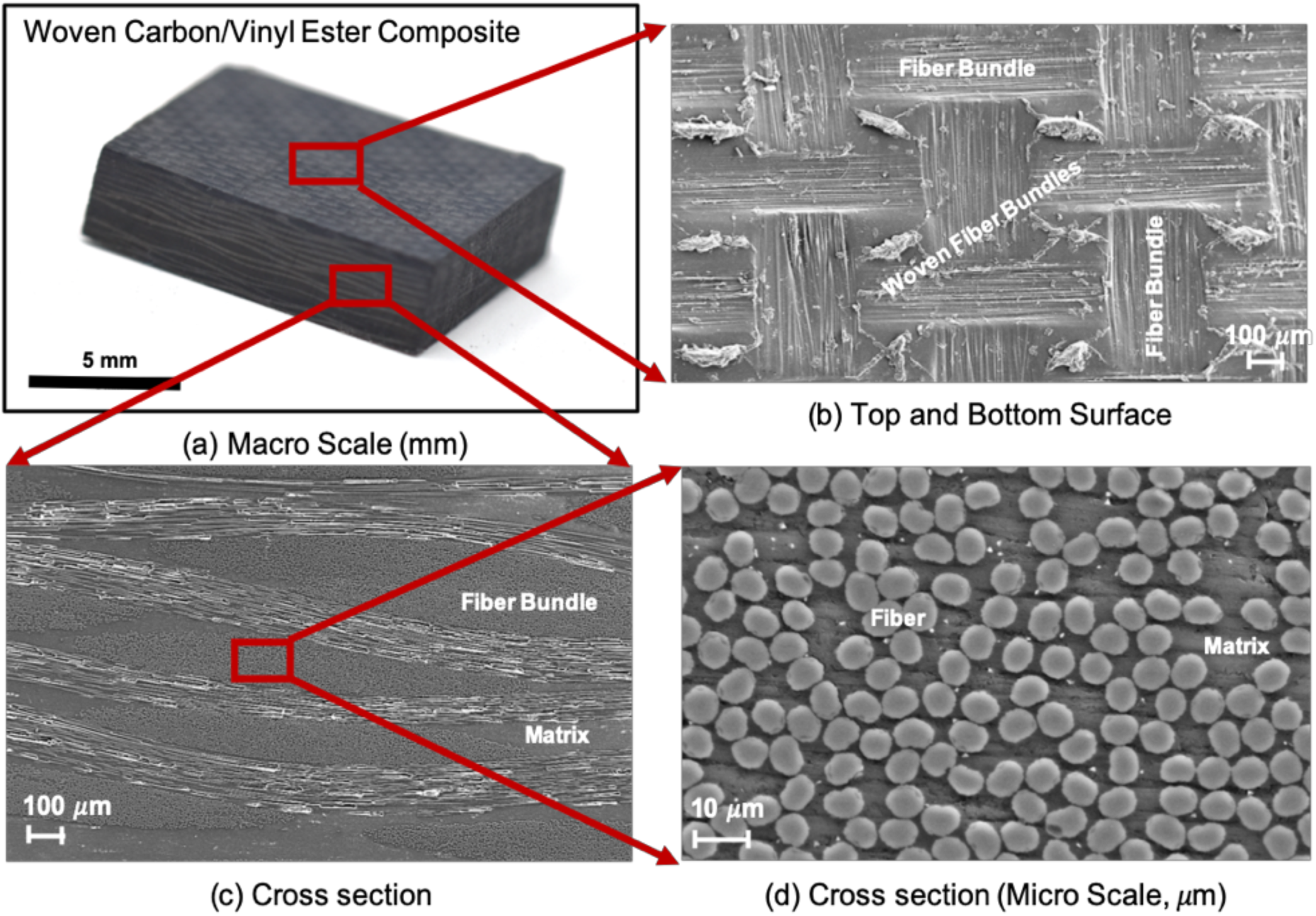
Optical and scanning electron microscopy (SEM) images of the fabricated polymer composite. Figure (a) shows an optical image of a polymer composite sample at the mm scale. This composite consists of woven bundles of carbon fiber with vinyl ester polymer as the binding matrix as shown in figure (b) and (c), which are SEM images of the top/bottom surface and cross-section of the sample in figure (a). Biofilms were extracted from top/bottom surfaces shown in figure (c). Figure (d) shows a cross-section SEM image within a fiber bundle which consists of fibers and matrix at the microscale.

**Figure 2.**
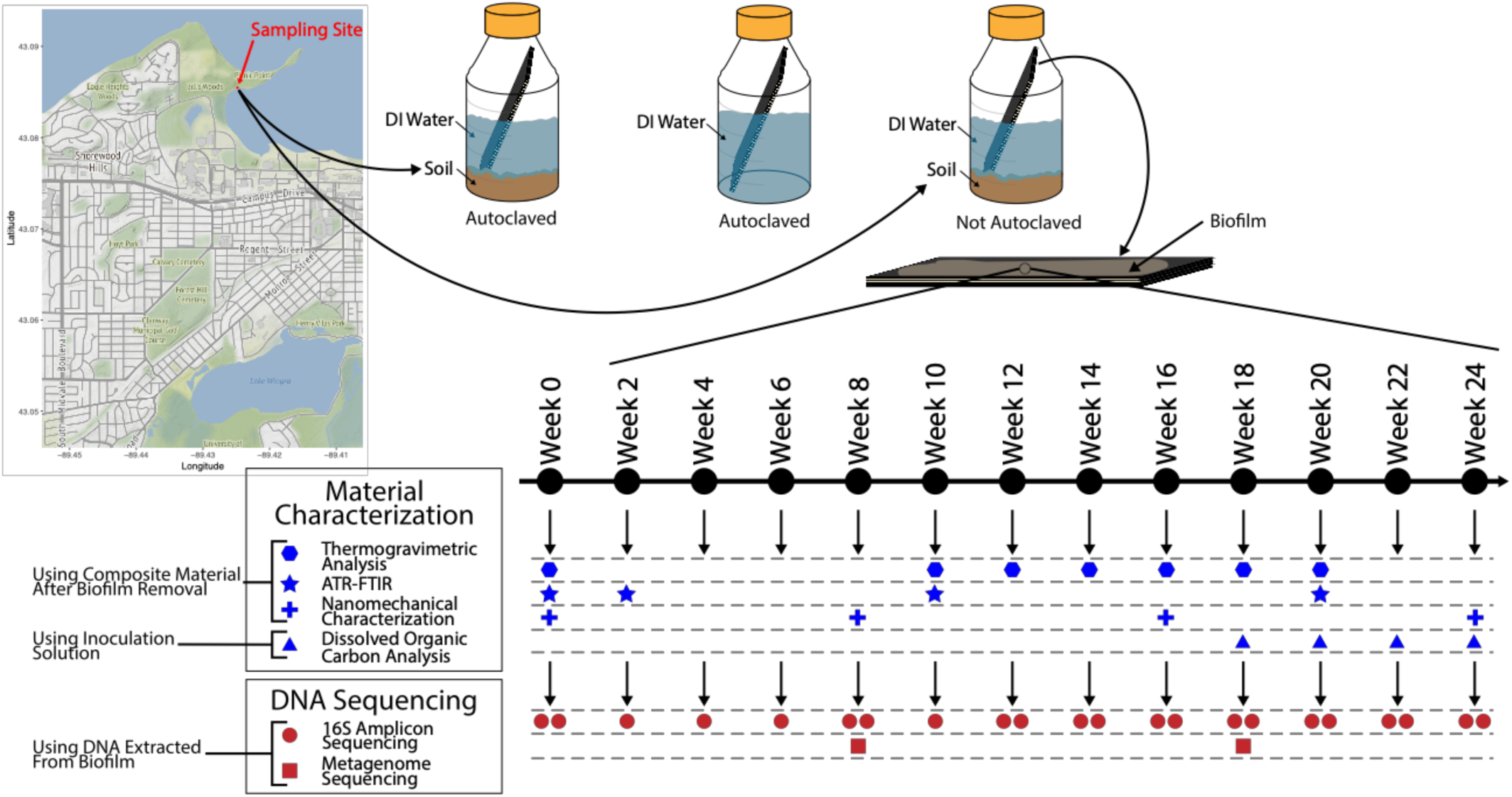
Sampling scheme for experimental study of degradation of polymer composites. Soil samples (collected from the location shown on the map) were used as inoculum for establishment of biofilms on polymer composites. Three different conditions were set up: S+DI+A, DI+A, and S+DI. All experiments were conducted in duplicate. The experiment was conducted over 24 weeks. Details of specific samples used for material characterization and DNA-sequencing based microbial analyses are shown. Three different materials characterization procedures were performed using Thermogravimetric Analysis, Fourier Transform Infrared Spectroscopy, and Nanomechanical Characterization. Dissolved organic carbon analysis was performed on liquid samples collected from the incubation solution for each week. Microbial analyses included 16S rRNA amplicon sequencing and genome-resolved metagenomics.

Three different materials characterization procedures, including TGA, FT-IR, and Nanomechanical Characterization, were performed on polymer composites. Possible leaching of carbon from the composites into the solution was measured using dissolved organic carbon analysis on liquid samples from the incubation solution towards the end of the tests. From these characterizations, we were able to clearly show that the manifestation of polymer composite degradation primarily resulted from microbial activity.

For microbial characterization, we performed 16S ribosomal RNA amplicon sequencing to determine the structure and abundance of the microbial community and metagenomic sequencing to profile the functional capacity of microorganisms to degrade polymer composites. To the best of our knowledge, no prior studies have investigated the combined effect of a diverse microbial community as occurs in nature on polymer composites. In contrast to previous studies that have focused on single isolates and their effects on composites, the vast majority of microorganisms in the polymer composite microbiome are uncultivated.

### Degradation of polymer composites in the presence of microbes

To establish the extent of degradation over varying exposure periods and to elucidate mechanisms of polymer degradation, we performed TGA upon exposure of polymer composites to different environments mentioned above. The extent of degradation in polymer composites was measured in terms of thermal decomposition onset temperature, which is expected to reduce with higher degrees of degradation. Typical output from a TGA is the weight loss percentage of a material sample when heated to high temperatures, typically in the range of 100 to 400°C which are plotted in a graph. The temperature at which the slope of this graph changes significantly (knee formation highlighted in Figure 3(a)) is referred to as the thermal decomposition onset temperature. We collected material samples from regions near the surface of polymer composite samples to conduct TGA. Figure 3(a) compares the TGA graphs of samples in soil solution (S+DI) from week 10 to week 20. We observed that the thermal decomposition onset temperature reduced with increasing exposure time. This was attributed to the hydrolysis of certain groups (e.g. ester groups, C=O groups) in the polymer chain that weakens their backbone structure^32^, resulting in polymer chain scission creating low molecular-mass compounds. Next, we compared the thermal decomposition onset temperatures for samples from week 10 to week 20 against exposure time for all environments in Figure 3(b). We observed that this onset temperature reduced significantly for polymer composite samples in soil solution (S+DI) as compared to autoclaved water (DI+A) and autoclaved soil with water (S+DI+A) conditions. We observed that the presence of microorganisms (in the [S+DI] samples) strongly correlated with a higher degree of degradation of polymer composites as compared to other conditions where no or minimal microbial growth and activity was possible. Hence, we conclude that microbial activity drives the polymer chain scission-based degradation of polymer composites, which was manifested as a substantial reduction of the onset melting temperature^33^.

**Figure 3.**
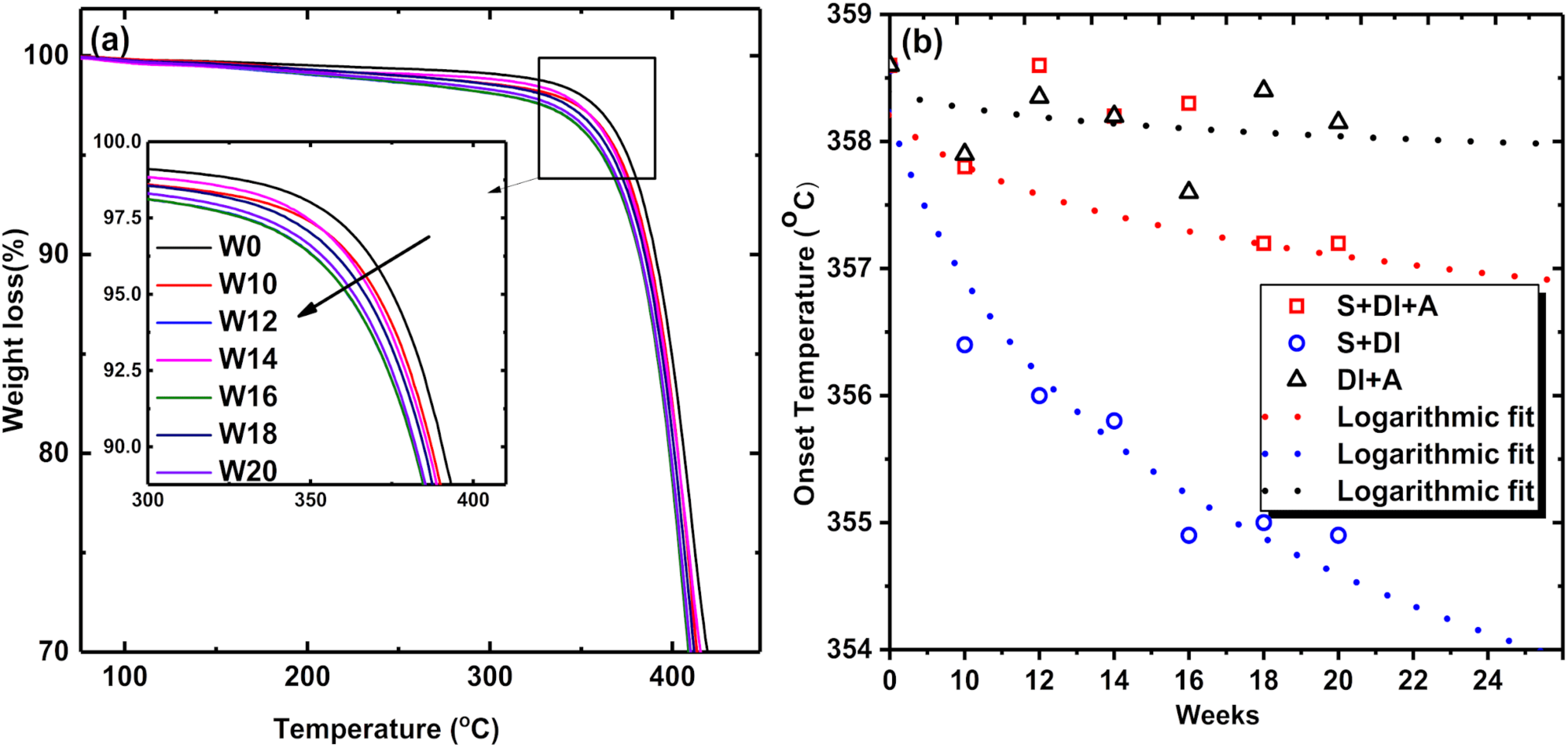
TGA of carbon fiber reinforced vinyl ester composites: **(a)** representative TGA weight loss (%) graphs with temperature for carbon fiber reinforced vinyl ester composites in soil with de-ionized water (S+DI) solution, and **(b)** graph of onset decomposition temperature (logarithmic fit) in different environments: soil with de-ionized water after autoclave (**□** - S+DI+A), soil with de-ionized water (O - S+DI), and de-ionized water after autoclave (**Δ**-DI+A), measured at different exposure time periods.

To establish the adverse influence of microbial degradation on the mechanical properties of polymer composites, we performed surface nanoindentation tests on soil solution exposed polymer composite samples. Reduction in mechanical properties due to degradation is measured in terms of modulus, hardness and displacement upon nanoindentation of polymer composites, where the reduction in modulus and hardness and increase in displacement are manifestations of polymer degradation. A comparison of load against displacement is a typical output from nanoindentation tests. Representative load - displacement graphs from nano-indentation tests on polymer composite samples exposed to soil solution (S+DI) from week 0, 8, 16 and 24 are shown in Figure 4(a). Modulus, hardness and displacement against the number of weeks were extracted from such graphs shown in Figure 4(a), which were then plotted in Figure 4(b). We observed that the modulus and hardness decreased with increasing number of weeks of exposure to soil solution, while the displacement increased. These results demonstrate that the surface nanomechanical properties of polymer composites were adversely affected by the degradation caused by microbial activity.

**Figure 4.**
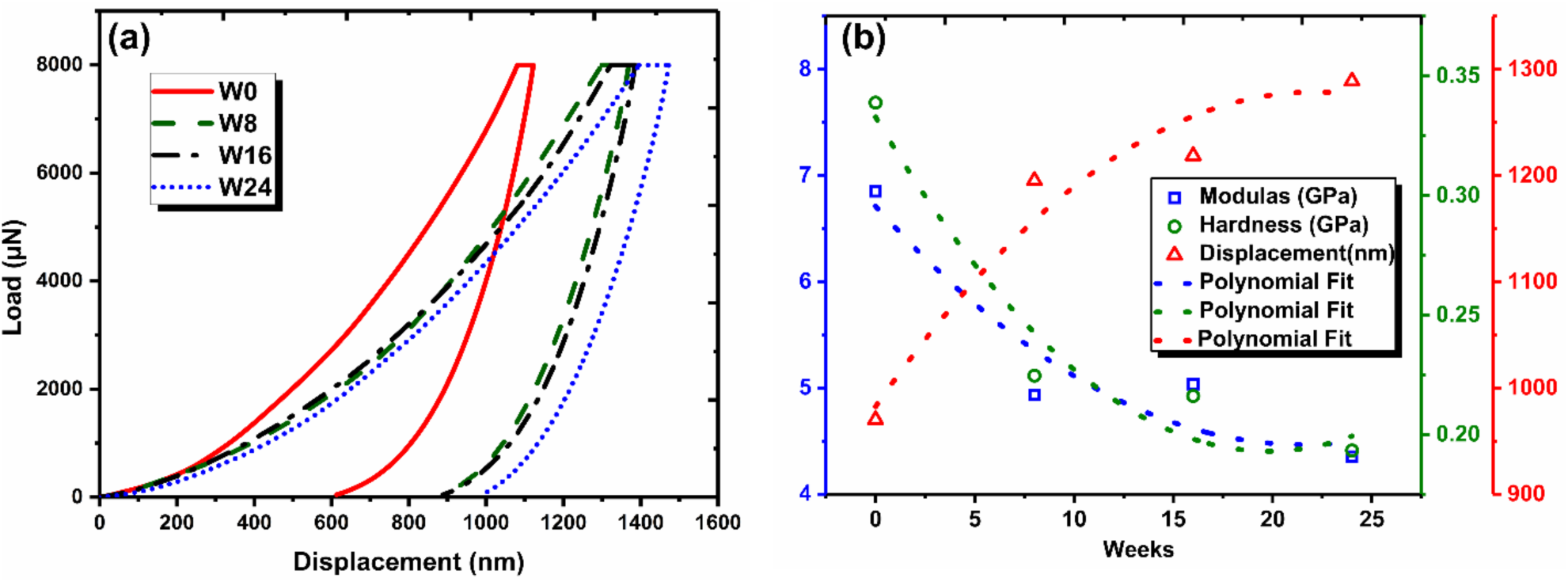
Nanomechanical characterization of carbon fiber reinforced vinyl ester composites in the resin-rich region near the surface: **(a**) load-displacement responses of carbon fiber reinforced vinyl ester composites after different exposure time periods in soil with de-ionized water (S+DI) solution, and **(b**) the mechanical responses such as Modulus (**□**), Hardness (O), and Displacement (Δ) with polynomial fit as function of different exposure time periods in soil with de-ionized water (S+DI) solution.

### Microbial diversity and abundance in the polymer composite microbiome

Given the presence of significant damage to polymer composites in the presence of soil, we hypothesized the involvement of microbial communities in their degradation. To determine the stability and diversity of the microbial community inhabiting the polymer composites microbiome, we collected biofilm samples every two weeks and used 16S rRNA amplicon sequencing to determine community composition. In total, we generated 2.2 million paired-end reads. Processing these reads revealed the presence of 8461 operational taxonomic units (OTUs) which is an estimate of the number of microbial genotypes in the polymer composite microbiome (Supplementary Table 1). Samples from weeks 0, 2, and 4 were distinct from other samples suggesting the recruitment of microbes into the biofilm from the soil samples used for inoculation (Supplementary Table 2, Supplementary Table 3). Microbial community composition and relative abundance across all sampling weeks are displayed in Figure 5. From weeks 6 to week 24, the microbial community inhabiting the polymer composite biofilm was distinct from the first three samples. In total, we found 446 OTUs to be unique to weeks 0, 2, and 4 while samples from weeks 6 to 24 had a total of 204 unique OTUs (Supplementary Table 2, Supplementary Table 3). The majority of the OTUs from weeks 6 to week 24 remained a stable component of the biofilm, but their abundance varied over the course of the 24 weeks. In total, only 8% of OTUs from week 0 were observed in week 24. Microorganisms from the bacterial lineages *Chlorobi, Deltaproteobacteria*, Candidate Phyla Radiation/*Patescibacteria*, and *Chloroflexi* were the most abundant members of the biofilm. To infer the metabolic capabilities of the dominant members of the biofilm to impact the polymer composite, we conducted metagenome sequencing on a subset of samples chosen from our microbial community composition analyses. We chose two samples from weeks 8 and 18 that represent the average microbial community across the 24 weeks of our experiment. Metagenome sequencing resulted in approximately 85.9 Gbp of paired-end reads. After read processing and assembly, we generated ∼100, 000 assembled DNA sequences (scaffolds) representing 1.74 Gbp of sequence (>1 Kb in length). To generate microbial genomes from assembled DNA sequences, we utilized a comprehensive genome binning and consolidation approach. This resulted in reconstruction of 144 metagenome-assembled genomes (MAGs) in total of which 51 MAGs were classified as high-quality and 70 MAGs as medium-quality drafts in accordance with MIMAG standards^34^. These 121 MAGs were used for downstream analyses of elucidation of metabolic potential and polymer composite degradation mechanisms of the biofilm. To undertake taxonomic classification of these genomes, we used two distinct phylogenetic reconstruction approaches including 16 concatenated ribosomal proteins and the Genome Taxonomy Database toolkit (GTDB-tk). Consistent with results from our 16S rRNA amplicon study, our phylogenetic analyses revealed that most MAGs represented the bacterial lineages *Chloroflexi* (17 MAGs), *Chlorobi* (33 MAGs), *Deltaproteobacteria* (24 MAGs), and Candidate Phyla Radiation (CPR)/Patescibacteria (25 MAGs) as shown in Figure 6 (a). Within the highly diverse CPR/Patescibacteria group, we recovered MAGs from the lineages Gracilibacteria, Moranbacteria, Saccharibacteria, Roizmanbacteria, CPR1, WWE3, Falkowbacteria and Yonathbacteria (Supplementary Table 4).

**Figure 5.**
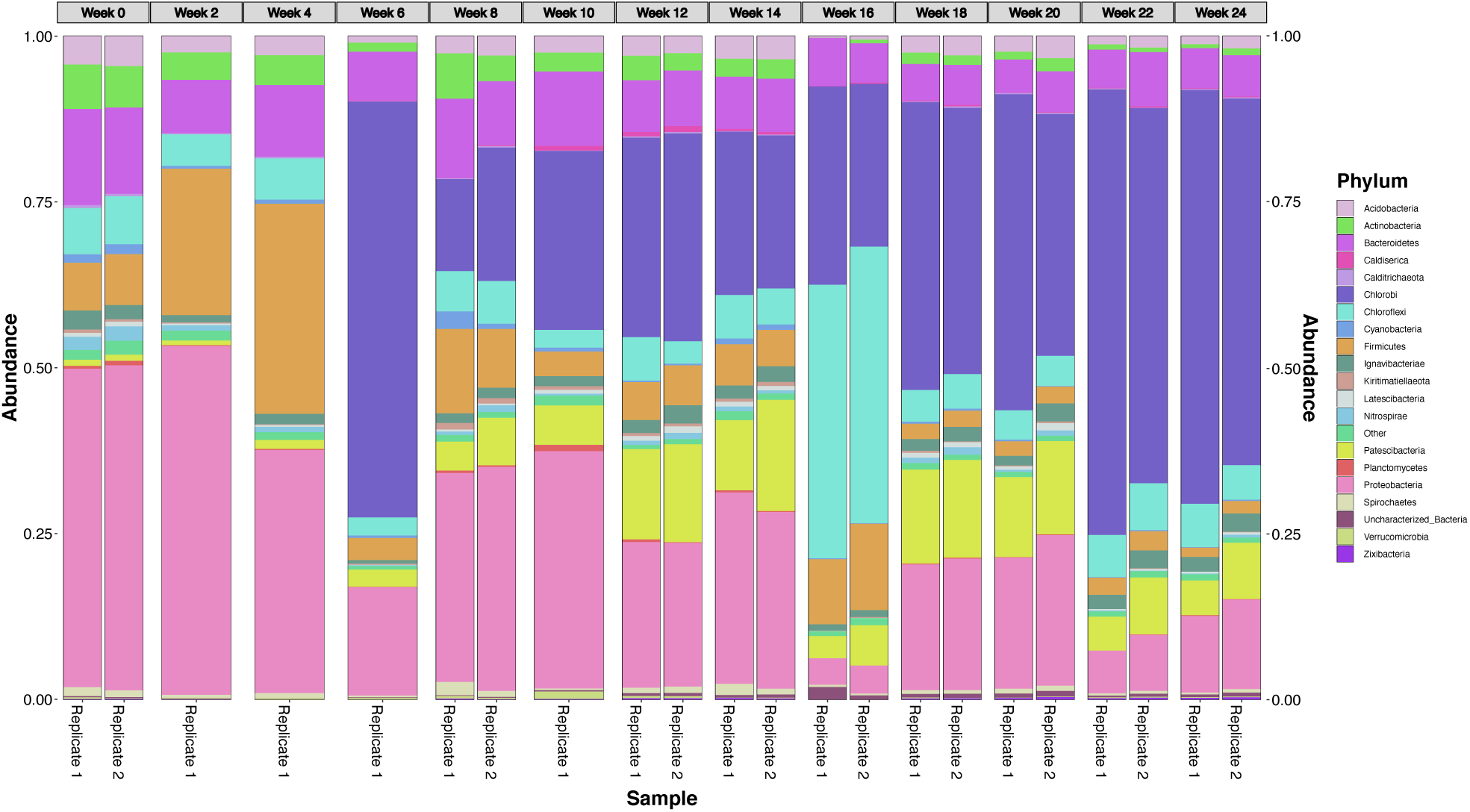
Microbial community composition and abundance measured using analyses of 16S rRNA Amplicons. Stacked bar graph describing the community composition of microbes growing on the carbon composite material across all samples. Weeks with two bars had duplicate DNA samples extracted and, therefore, had duplicate 16S community abundance plots generated for them.

**Figure 6.**
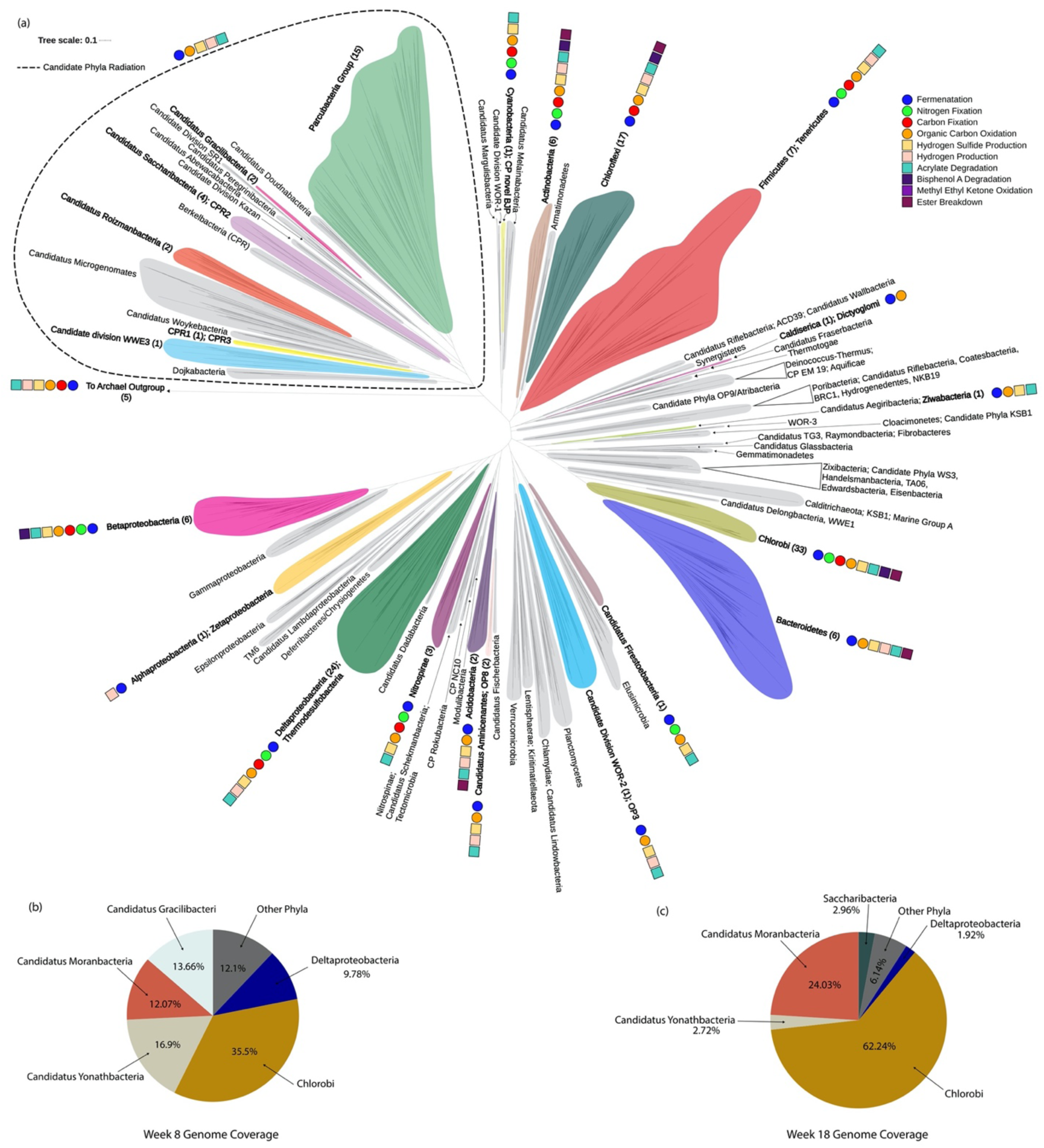
Phylogenetic and Relative Abundance Analysis of organisms in the polymer composite biofilm. (**a**) Phylogenetic tree created by alignment of 16 ribosomal proteins of metagenomes curated from two different weeks of sampling. Colored gray are those Phyla and groups of Phyla that did not contain representatives from our set of metagenomes. For Phyla where metagenome sequencing resulted in the presence of genomes within those Phyla, branches are surrounded by colors other than gray and Phyla names are in bold. Indicated with a dashed line are all genomes used to create this tree that fall within the group, Candidate Phyla Radiation. **(b)** Pie chart describing the percentage of the total coverage values generated from the week 8 genomes associated with each specific Phylum listed. Phyla responsible for the lowest percentages of the total coverage values were grouped into “Other.” **(c)** Pie chart describing the percentage of the total coverage values generated for the week 18 MAGs associated with each Phylum listed. Phyla responsible for the lowest percentages of the total coverage values were grouped into “Other.”

To estimate the relative abundance of all organisms in the biofilm, we conducted read-mapping to estimate the coverage of individual MAGs across weeks 8 and 18. Our results show that MAGs from four bacterial lineages *Chloroflexi, Chlorobi, Deltaproteobacteria*, and CPR/Patescibacteria accounted for 91 - 95% of the total microbial abundance (Figure 6 (b), Figure 6 (c), Supplementary Table 5). Specifically, organisms from the CPR/Patescibacteria group comprised 30 - 43% of the abundance. Nearly, the entire abundance of CPR/Patescibacteria could be attributed to organisms from the lineages, *Candidatus* Gracilibacteria, *Candidatus* Moranbacteria, *Candidatus* Yonathbacteria, and *Candidatus* Saccharibacteria.

### Metabolic potential of the polymer composite microbiome

The 121 high- and medium-quality MAGs from the polymer composite microbiome encoded a total of 275,719 protein-coding open reading frames. To determine important metabolisms encoded by the abundant organisms in the biofilm, we screened these proteins for their association with metabolic pathways. This approach served two key purposes. First, the association of dominant organisms with metabolic pathways sheds light on potential biological processes that occur in polymer composite biofilms in nature. Second, this allows us to identify specific metabolic pathways that are associated with the breakdown of individual components in polymer composites. Importantly, we identified energy metabolism and key ecosystem functions being conducted by organisms in the biofilm that can shed light on their roles, and their importance to degradation of polymer composites. Metabolisms encoded in the polymer composite microbiome included sulfate reduction, sulfur oxidation, fermentation, nitrogen fixation, carbon fixation, and organic carbon oxidation (Supplementary Table 6, Supplementary Table 7, Supplementary Table 8, Supplementary Table 9). The metabolic potential for sulfate reduction was observed in *Deltaproteobacteria* and *Nitrospirae*. Sulfur oxidation was observed in *Chlorobi* and *Betaproteobacteria*. In accordance with previous observations, all *Chlorobi* organisms were phototrophs also conducting sulfide oxidation. Fermentation, primarily acetogenesis was observed in organisms from 18 different phyla including CPR. While other organisms from other phyla complemented their metabolism with fermentation, CPR organisms were not observed to possess any alternate forms of metabolism (Supplementary Table 6, Supplementary Table 7). The ability to fix nitrogen was observed in 32 MAGs from 10 distinct lineages. Finally, carbon fixation was a commonly observed trait in the biofilm. Organisms from ten different phyla possessed different pathways for carbon fixation including the reverse tricarboxylic acid (TCA) cycle and the Wood– Ljungdahl pathway. To estimate the connected nature of microorganisms and metabolism in the composite microbiome, we estimated energy flow (Supplementary Figure 1) and metabolic connections (Supplementary Figure 2) in the microbial community. Our results show that the composite biofilm was not dominated by a single metabolism but was instead highly connected and diverse metabolisms associated with carbon, nitrogen and sulfur transformations contributed to energy metabolism.

### Structural chemical composition changes in polymer composites due to microbial activity

To understand the structural chemical changes introduced by microbial activity in polymer composites, we performed a detailed surface chemical analysis using FT-IR (Supplementary Table 10, Supplementary Table 11, Supplementary Table 12). A typical output of FT-IR is a comparison of absorbance intensity with respect to wavenumber (frequency) (Figure 7(a), Supplementary Figure 3, Supplementary Figure 4), which is a result of the interaction of infrared radiation between the chemical bonds at different frequencies. The structural chemical changes before and after exposure to different conditions were analyzed. Figure 7 (a) shows superimposed FT-IR spectra of a polymer composite sample exposed to soil solution for different exposure times. The spectra show several peaks corresponding to different wavenumbers that manifest due to the interaction of infrared radiation between chemical bonds at different frequencies. Changes in chemical bonds are reflected as changes in peak intensity, broadening, and shifting with respect to the peaks expected at specific frequencies corresponding to a particular type of chemical bond. Peak broadening and higher peak intensities are associated with increased formation of intramolecular or inter-molecular hydrogen bonding^35^.

**Figure 7.**
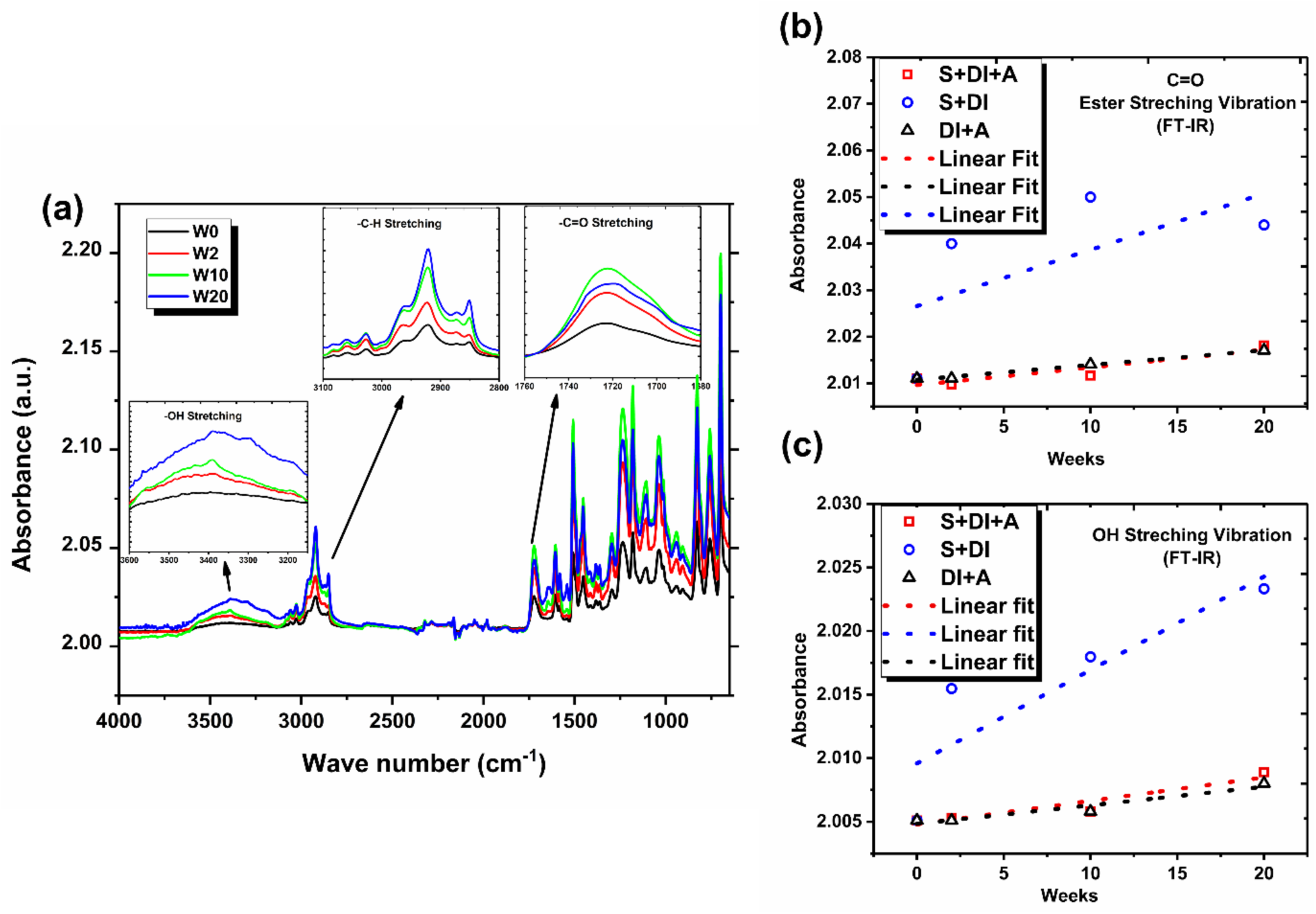
Attenuated total reflection Fourier-transform infrared spectroscopy (ATR-FTIR) analysis of carbon fiber reinforced vinyl ester composites: **(a)** superimposed ATR-FTIR spectra of carbon fiber reinforced vinyl ester composites in soil with de-ionized water (S+DI) solution after different exposure times confirm peak broadening and increase in peak intensity. The characteristic peaks corresponding to -OH stretching (moisture), -C-H stretching (vinyl), and -C=O stretching (ester) are shown in the insets of Figure (a). Peak intensity increased and broadened for the characteristic peak of vinyl ester corresponding to C=O stretching vibration (1726 cm^-1^)^72^ with increasing exposure time (from week 0 to week 20) when samples were exposed to microorganisms (S+DI samples). Similar increases observed in peaks corresponding to aromatic and aliphatic C-H stretching (2800-3100 cm^-1^) and O-H stretching (3200-3600 cm^-1^)^72^. Increase in absorbance intensities with exposure time corresponding to **(b)** -C=O stretching vibration, and **(c)** -OH stretching of carbon fiber reinforced vinyl ester composites with linear fit after exposing them to different conditions: soil with de-ionized water after autoclave (**□**-S+DI+A), soil with de-ionized water (**□-**S+DI), and de-ionized water after autoclave(**Δ**-DI+A).

In Figure 7 (b) and (c), C=O stretching and -OH stretching absorbance intensities were summarized for samples exposed to different conditions over the period of 20 weeks. The absorbance intensities increased with exposure time when compared to samples from week 0, indicating a change in the chemical molecular structure. In particular, this change was significantly higher for samples in the presence of microbial activity as compared to where microorganisms were absent, or in controls.

To further elucidate the impact of microbial action on the degradation of polymer composites, we analyzed the dissolved organic carbon (DOC) content in each of the solutions after conditioning the composite samples. In general, the amount of DOC content in the solution should increase based on the chain scission mechanism associated with hydrolysis of esters^36^. Here, we performed DOC analysis to support and confirm the chain scission mechanism described in Figure 8 (a). Water molecules preferentially establish a weak hydrogen bond with the vinyl ester polymer, which manifested as peak broadening of the -OH stretching from FT-IR analyses. Weak hydrogen bonding promotes the chain scission process over time, which causes lower surface nanomechanical properties such as lower hardness and modulus, which was confirmed by nanomechanical characterizations in Figure 4.

**Figure 8.**
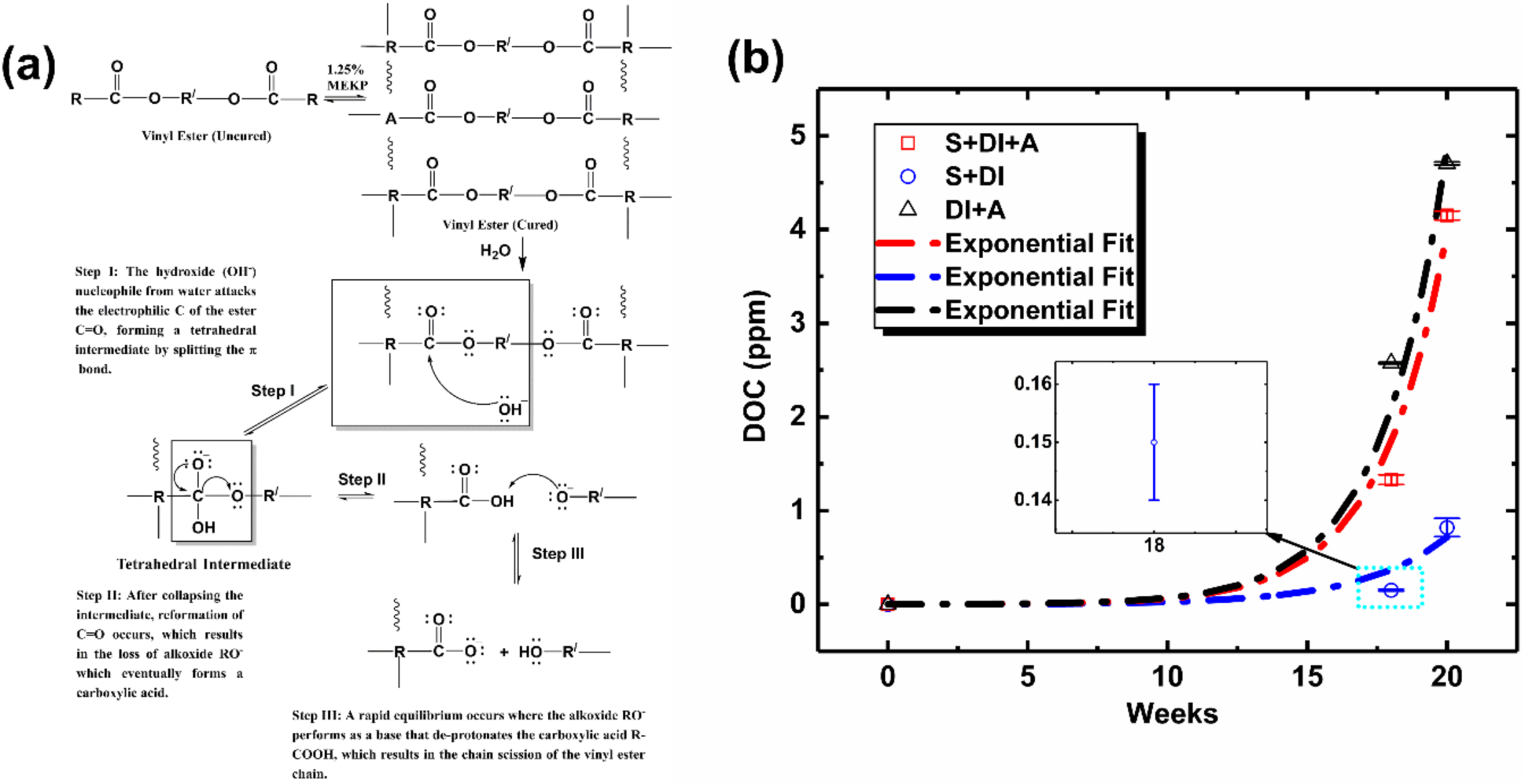
Chemical structural alteration of vinyl ester resin after exposing to different conditions at different exposure due to polymer chain scission: **(a)** mechanism explained - hydrolysis process of carbonyl group causes chain scission of esters, which results in dissolved organic carbon (DOC) content in the solution, **(b)** change in the amount of DOC content (Supplementary Table 19) (exponential fit) for different conditions: soil with de-ionized water after autoclave (**□**-S+DI+A), soil with de-ionized water (O-S+DI), and de-ionized water after autoclave (**Δ**-DI+A).

As seen in Figure 8 (b), the DOC content increased with increasing exposure time for composite samples under different conditions. However, the increase in DOC in soil solution with time is not as pronounced as compared to other conditions. This is attributed to the presence of microbes that consume/alter the DOC content over time^37^, and a cyclic recovery of DOC is needed to maintain equilibrium in the solution. This phenomenon results in a cyclic process of further chain scission, DOC consumption/alteration, and microbial growth, which causes severe degradation of the polymer composites. In other cases without microbes, DOC content increases significantly as there are no microbes present to consume or alter the DOC content. In these cases, the chain scission process will stop in order to maintain equilibrium, and the mechanism proposed in Figure 8 (a) will not continue to degrade the polymer composite.

### Potential contributions of microbial metabolism to degradation of polymer composites

To understand the microbial mechanisms driving the degradation of polymer composites, we identified metabolic pathways in microbial genomes that can transform or breakdown individual components of polymer composites. In total, we identified six unique metabolisms that had detrimental effects on the structure of the polymer composite by causing breakup of the physical structure through gas formation and blistering, and degradation of organic compounds in the matrix through chain scission and other processes^24,28^ (Table 1, Supplementary Table 13). Prior research has suggested that production of gases such as hydrogen and hydrogen sulfide gases can degrade the binding matrix and decrease the structural integrity of the composite^24,28^. The metabolic capability to produce hydrogen was identified in 24 MAGs from 11 different lineages involving three hydrogen-evolving hydrogenases, including two families of [FeFe] hydrogenase and a group 4 [NiFe] hydrogenase^38^. Amongst the four most abundant lineages in the composite microbiome, we detected this capacity in MAGs from CPR/Patescibacteria, *Deltaproteobacteria* and *Chloroflexi*. Two distinct microbial pathways can produce hydrogen sulfide by the anaerobic respiration of sulfate/sulfite^39^, or by degradation of the amino acid cysteine^40^. While both of these metabolic capacities were detected in the composite microbiome, degradation of cysteine was observed to be more abundant. The ability to respire sulfate/sulfite and produce hydrogen sulfide was observed only in MAGs from the lineages *Deltaproteobacteria* and *Nitrospirae*.

**Table 1.** Key microbial mechanisms associated with degradation of polymer composites.

| Mechanism of polymer composite degradation | Potential microbial enzymes/proteins involved |
| --- | --- |
| Hydrogen production | Ni-Fe Hydrogenases, Fe-Fe Hydrogenases |
| Hydrogen sulfide production | Sulfite reductases, Cysteine metabolism |
| Acrylate degradation | Amidases |
| Bisphenol A degradation | Bisphenol A hydroxylase |
| Methyl ethyl ketone oxidation | Butanone monooxygenase |
| Breakdown of esters | Esterases |

We identified four components of the polymer composite binding matrix as capable of being degraded by microbial action, including methyl acrylate, bisphenol A, methyl ethyl ketone, and esters. Amongst these, the most abundant capacity in the composite microbiome was the ability to degrade acrylate which was observed in 78% of all MAGs (Supplementary Table 14). The ability to degrade bisphenol A and esters were less common in the biofilm and identified in 7 MAGs each. No pathways for the degradation of methyl ethyl ketone were detected in any of the organisms. Since we did not directly measure the activity of these processes, we estimated growth rates of microorganisms likely to be associated with degradation of composites. This was performed using a recently demonstrated approach where it was shown that growth rates of microbial strains in their natural environment can be determined by measuring the ratio of the coverage of DNA at the origin and terminus of replication in a genome^41^. We observed that all four abundant lineages, *Chlorobi, Chloroflexi, Deltaproteobacteria*, and the CPR/Patescibacteria were growing actively. We propose that a significant proportion of these organisms likely source carbon from the polymer composites and their high rates of growth are associated with breakdown of the polymer composite (Supplementary Figure 5).

## Discussion

Polymer composites are being used increasingly in lightweight structural applications in the built environment and will likely be a backbone of next-generation materials. Therefore, significant resources and research are needed to understand the performance and durability of these materials under physical, chemical, and biological stressors that can induce damage and impact their durability. Our goal of this study was to elucidate biologically driven degradation mechanisms of polymer composites. To that end, we studied the degradation of polymer composites in the presence of microbial communities, and identified key modes of biologically driven degradation which are likely associated with selective microbial enzymes.

Our temporally resolved experimental design involved inoculation of polymer composites with microbial communities from natural soils. We recorded the early onset and growth of a biofilm on the surface of these polymer composites. Overall, we observed a significant reduction in the molecular weight of the vinyl ester resin used as a binder in polymer composites. With increasing exposure time to microbes, a greater reduction in molecular weight was observed. We propose this to be a manifestation of chain scission in the vinyl ester resin of the polymer composites. Vinyl ester resins are composed of a number of organic compounds, of which acrylate, bisphenol and esters are the most abundant. Specifically, we observed that certain bonds such as C=O, aromatic and aliphatic C-H, O-H and -OH, which are common in the vinyl ester resin were specifically targeted by microbial activity. We identified four microbial groups to be especially abundant components of the microbial biofilm, namely CPR/Patescibacteria, *Deltaproteobacteria, Chlorobi*, and *Chloroflexi* and contributed towards acrylate degradation, bisphenol A degradation, breakdown of esters, hydrogen production and hydrogen sulfide production.

Amongst the microbial metabolisms associated with degradation of the binding matrix, the most abundant capacities in the composite microbiome were associated with degradation of acrylate, and production of H_2_S (Figure 9). The ability to degrade bisphenol A and esters, and produce hydrogen were less common in the biofilm. We speculate that microorganisms that can degrade acrylate and/or produce H_2_S are selected for in this environment. There are three potential factors that support these observations. First, acrylate degradation is primarily mediated by amidases that are abundant in these specific microbial groups. While amidases are primarily associated with degradation of amide containing compounds such as urea for sourcing nitrogen, we propose that the composite microbiome utilizes these for degradation of acrylate in the vinyl ester resin^42^. Second, H_2_S may be produced by dissimilatory sulfate reduction by organisms from the *Deltaproteobacteria* and *Nitrospirae* lineages^39^. Prior research has demonstrated gradients of oxygen in biofilms including the presence of anoxic regions that could support highly active sulfate reducing microbes^43^. Third, H_2_S may also be produced by degradation of cysteine which is likely an abundant component of organic-rich biofilms^40^. We propose that the interplay between the degradation of the binding matrix (acrylate, bisphenol A, esters) through chain scission of chemical bonds and the production of gases (H_2_S, H_2_) that can cause physical damage and potential bloating of the polymer structure contributes towards accelerated degradation of polymer composites.

**Figure 9.**
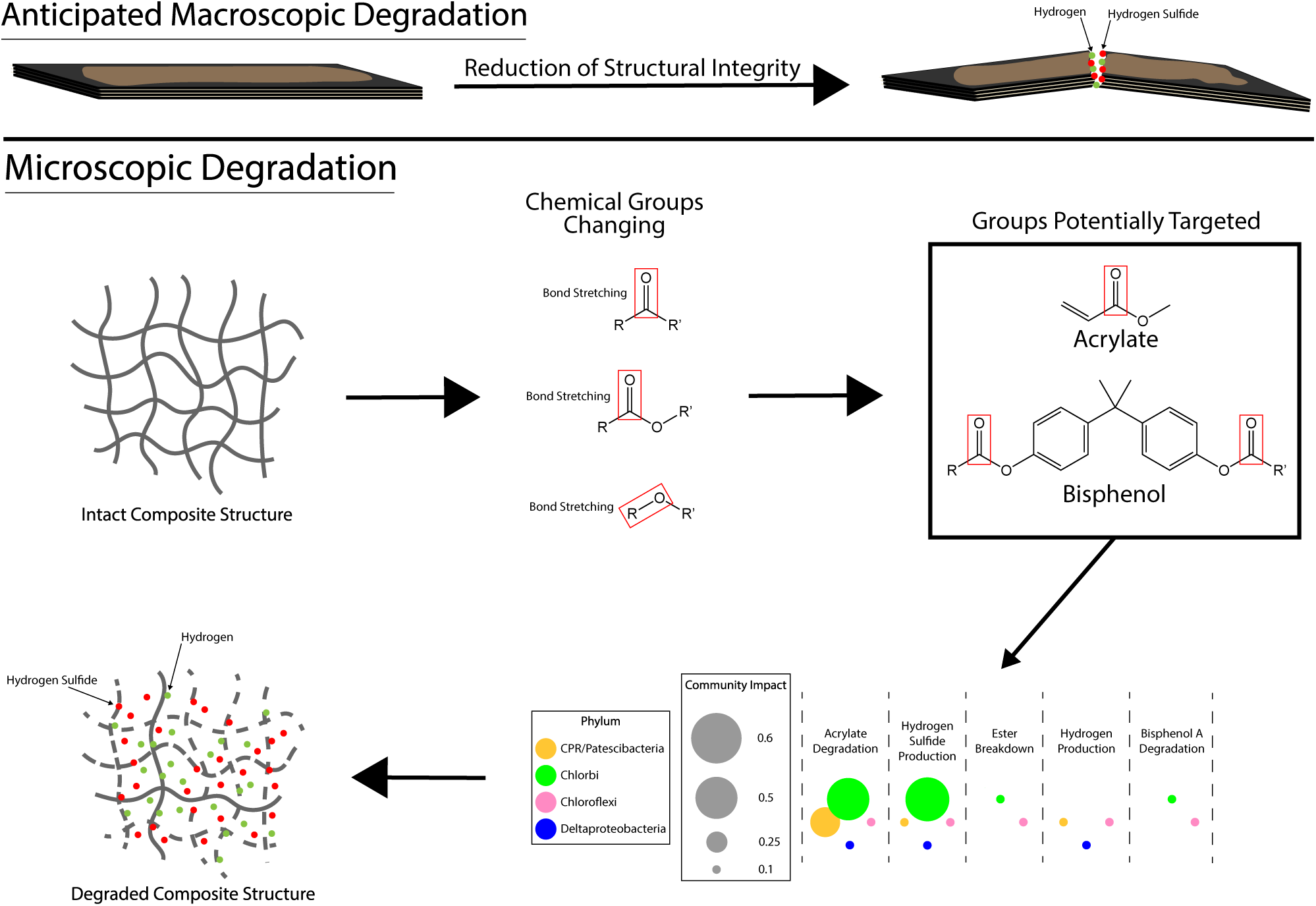
Macroscopic and microscopic degradation of polymer composite structure. Macroscopic degradation, which manifests in reduction of structural integrity, is shown as the progression from an intact composite to an exaggeratedly broken composite. Microscopic degradation manifests itself as breaks in the polymer chain, which is influenced my microbial metabolisms. Chemical groups being modified were identified through FT-IR and were extrapolated to groups within compounds present in the composite material.

Our study builds on prior research involving studies of single microorganisms such as *Pseudomonas, Lactococcus*, SRB, *Rhodococcus*, and *Ochrobactrum* species^21,24,44^ that can degrade polymer composites. Identification of microorganisms from four bacterial phyla, namely, *Deltaproteobacteria, Chlorobi, Chloroflexi*, and CPR/Patescibacteria as dominant components of the microbial biofilm on polymer composites provides vital insights that could direct approaches for testing the durability of these structures in the field. Organisms from these bacterial lineages are abundant in a variety of terrestrial, marine and coastal environments where they could degrade polymer composite-based infrastructure^45,46^. In remote field applications, such as in pipelines and wind turbines, any observed biofilms on the composites could be tested for the presence of the above microbial lineages and could indicate ongoing deleterious effects and help mitigate damage. While our study focused on specific modes of degradation of polymer composites, it is possible that additional currently unknown mechanisms exist for this process. Future studies should complement genome-based analyses with measurements of activity and isotope-based quantification of the magnitude of degradation of polymer composites.

A recent report describing the state of infrastructure in the U.S. from the American Society for Civil Engineers revealed that current infrastructure in the U.S. is deteriorating rapidly, and failure to act towards improving these would be an issue for the nation’s public safety, economy, and international competitiveness^47^. Fiber-reinforced polymer composite products and systems can provide cost-effective replacements for rehabilitating or retrofitting existing infrastructure compared to conventional construction materials used in harsh environments. For example, polymer composites can be used for rehabilitating structures made of steel, timber or concrete at about 5% of their replacement cost. In recent years, there has been an increasing demand for using polymer composites in designing new or rehabilitating existing infrastructure under harsh environments. Given the lightweight nature of polymer composites, they are also excellent candidates for applications in remote regions, for example, like the Arctic. As the Arctic region evolves due to the melting of sea ice, we will encounter novel physical, chemical, and biological factors that are not prevalent in regular environments, and an approach like the one developed here can assist in identifying and modeling durability of polymer composites in the Arctic.

Microbial activity is an important environmental factor in most marine, terrestrial, and coastal environments, where these polymer composites are likely to be used. The five important microbially-mediated degradation mechanisms of polymer composites recognized in this study motivates new thinking about how their durability should be modelled. Although the current study focuses on polymer composites with vinyl ester resin, the observations and conclusions are also relevant and beneficial to other resin systems like epoxies and polycarbonates, where acrylate and bisphenol are key compounds. Hence, we propose that microbial interactions with polymer composites should be considered in future modeling of polymer composites developed for long-term durability and life prediction.

## Methods

### Thermogravimetric Analysis

TGA is a thermal analysis tool used for evaluating the physical and chemical changes in materials as a function of temperature. A material sample is heated at a constant heating rate and the temperature-dependent mass change is established. Samples collected from different environments subjected to TGA usually present temperature-dependent mass change, which is related to the chemical or physical changes due to temperature and specific environmental conditions. Herein, we conducted TGA on polymer composite samples using a Netzsch TG 209 F1 Libra (Netzsch Instruments North America, LLC) setup fit with nitrogen as the purge gas. We used nitrogen gas to maintain an inert atmosphere in the test chamber to act as a protective gas for our sample and instrumentation. We collected samples from the region close to the surface of the polymer composites, which were periodically taken from different conditions. We prepared three samples for each condition to obtain reproducible thermal properties. We maintained a similar thermal mass for each sample, which was around 15-20 mg of material. Netzsch TG 209 F1 Libra is designed for 0.1 μg TGA resolution, which means it can precisely measure the change of mass up to 0.1 μg at any given heating rate and temperature. After collecting a sample from a polymer composite using fine sharp scissors, we placed it in an 85 μl alumina sample pan (outer diameter of 6.8mm x height x 4mm). We used an alumina pan to avoid any contamination from the pan. After that, we heated the sample from room temperature to 600 °C at a heating rate of 5 °C/min with a 40 ml min^-1^ flow rate of nitrogen. This process was repeated for every sample collected from the polymer composites exposed to different conditions.

### Chemical structure analysis

FT-IR is a chemical analysis tool specially designed for organic materials in order to verify their chemical structure. This analysis uses the principle of the interaction of infrared radiation between the chemical bonds at different frequencies. This interaction induces vibrational excitation of covalently bonded atoms and groups that have a particular energy intensity at a certain frequency. A typical output of FT-IR is a graph of absorbance /transmittance intensity with respect to wavenumber (frequency). A standard FT-IR spectral library is available for all possible chemical bonds that serves as a fingerprint to match a measured infrared radiation (IR) spectra. For example, C=O bonds interact with the IR frequency in a range of 1650 cm^-1^ - 1800 cm^-1^, which manifests as absorbance/transmittance peak at this frequency range. Changes in chemical bonds are reflected as changes in peak intensity, peak broadening, and peak shifting with respect to the peaks expected at specific frequencies corresponding to a particular type of chemical bonding. In this study, we performed FT-IR to observe changes in the chemical structure of polymer composites before and after exposure to different conditions. We used Bruker Tensor 27 (Thermo Scientific Instrument) equipped with attenuated total reflectance (ATR) to analyze the chemical structures of the composites. This instrument is also equipped with a room temperature deuterated triglycine sulfate (DTGS) detector, mid-IR source (4000 to 400 cm^-1^) and a KBr beam splitter. This instrument has the capability of scanning IR in the range of 4000-400 cm^-1^ at an increment of 1 cm^-1^, which indicates the degree of fineness of the data obtained by the measurements. Before we placed our sample on ATR sample cell, we ran a background scan with the empty cell using 32 scans at 4 cm^-1^ resolution within a range of 4000-650 cm^-1^ to obtain a baseline. We chose the number of scans and resolutions to obtain a smooth spectrum, and used the mid -IR source (4000 to 400 cm^-1^) to obtain the IR spectrum. We chose a range of 4000-650 cm^-1^ to obtain the IR spectrum as it encompasses all the anticipated chemical bonds of interest in this study. After that, we placed the square-cut specimen on top of the ATR sample cell equipped with a single diamond crystal. After placing the sample on top of the crystal spot, the arm rotated over and turned down to press the sample down onto the diamond crystal face to get better contact. Then, we collected the data using Bruker OPUS Data Collection software using 32 scans at 4 cm^-1^ resolution within the range of 4000-650 cm^-1^. We chose the instrument’s default wide-open aperture settings (∼6 mm) to obtain maximum infrared interaction with our specimen. The aperture is placed in the line of IR beam between the IR sources and interferometer.

### Nano-mechanical measurements

Nanoindentation is a mechanical measurement tool that enables us to measure mechanical properties such as modulus and hardness in nanoscale. Primarily, load and depth of indentations are measured during the experiments. Different shapes of indenters have the ability to measure different nanomechanical responses, for example, Berkovich type indenter is used for modulus and hardness, cube corner for fracture toughness, spherical cone for scratch and wedge for three-point bending (flexural) measurements. In this study, we performed nanomechanical measurements in a Hysitron triboindenter TI900 equipped with an optical microscope for imaging. We chose Berkovich type Nano indenter for nanoindentation, which was calibrated using a reference specimen of fused silica and acrylonitrile butadiene styrene (ABS). We performed the nano-mechanical tests as per ASTM standard E2546 - 15^48^ on polished cross-sections of composite specimens, which were periodically collected from the soil solution condition. For mechanically grinding and polishing the polymer composite cross-section, we used 400-1200 grit SiC metallographic abrasive grinding paper (Pace Technologies, USA) for fine grind and then polished with 3μm diamond solution on DACRON II Polishing Cloth (Pace Technologies, USA). Upon polishing, we mounted the specimens on a magnetic disk using epoxy glue and then performed nanoindentation on the specimens. We applied a maximum load of 8000 μN (*P*_*max*_) at a loading rate of 1600 μN/s. This maximum load was maintained for 2 sec, after which the specimen was unloaded at the same rate of 1600 μN/s. We performed a minimum of 10 indentations on each assessed region for reproducibility. We used the unloading graph of each indentation to obtain the hardness and modulus value using equation 1-3.

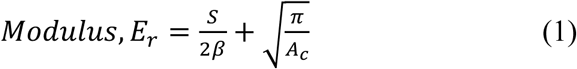

where S = Contact stiffness (*dp/dh*), *β = 1* for a Berkovich indenter with the angle of 65.27°, *A*_*c*_ = projected contact area at *h*_*max*_, *h*_*max*_*= maximum depth (displacement) corresponding to P*_*max*_

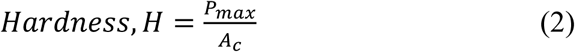

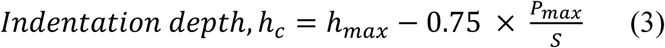

### Dissolved organic carbon

Dissolved organic carbon (DOC) is defined as the fraction of total organic content (TOC) in a solution that passes through a 0.45 μm filter. The part remaining on the filter is called particulate organic carbon or non-dissolved organic carbon (POC/NDOC). We collected solutions from different conditions used in this study and filtered them individually through a 0.45μm pore size filter (Nalgene, 045um syringe filter, 25-mm cellulose acetate membrane). We then transferred the solutions into centrifuge tubes/vials (PP graduated, 50 ml). We centrifuged (Eppendorf 5804R centrifuge) the solutions at a rate of 4500 rpm for 60 seconds and decanted the top solution after that. We then transferred the filtered and decanted solution to a vial. The vials were baked at 400°C for sterilization before using them for measurements. We kept the vials serially on the sample holder in addition to three blank vials for measurements with deionized water (electrical resistance - 18MΩ) to obtain a baseline. We then programmed the procedure for each measurement in a GE Sievers autosampler software, where we chose the default reagent process for blank run and chose auto reagent process for unknown runs. We flushed the system (GE Sievers M5310C, USA) with deionized water (electrical resistance −18MΩ) three times before obtaining each measurement. We measured the DOC concentration in solutions (20ml) from different conditions using a TOC analyzer (GE Sievers M5310C, USA), and observed that the measurement error was within ±3%.

### Collection of Microbial Samples

Microbial biofilms were collected carefully using sterile techniques from the polymer composite surface and stored frozen for subsequent DNA extraction. Biofilm sampling was performed once every two weeks for 24 weeks, with duplicate samples being taken on all samples, except for weeks 2, 4, and 10, due to inadequate growth of the biofilm. Samples were stored at −80°C until DNA was extracted, or the samples were otherwise utilized.

### DNA Extraction

DNA was extracted from biofilm samples using a modified version of the protocol from the Qiagen DNeasy PowerSoil Kit (Qiagen, Hilden, Germany). To modify the protocol, homogenization by vortexing was replaced by homogenization with the MP Biomedicals FastPrep-24 5G machine. Two samples whose concentrations were not sufficient for sequencing were concentrated by spinning in a Savant SpeedVac SC100 for 30 minutes on medium heat. 16S rRNA Amplicon sequencing was performed at the University of Wisconsin—Madison Biotechnology Center DNA Sequencing Facility (Madison, WI, USA) on the Illumina MiSeq instrument (Illumina, Inc., San Diego, CA), producing 2×300 bp reads. 16S amplicon sequencing was performed on all DNA samples obtained in the study (Weeks 0, 2, 4, 6, 8, 10, 12, 14, 16, 18, 20, 22, 24). Replicate DNA samples were obtained and 16S Amplicons were sequenced for all weeks except Weeks 2, 4 and 10. Metagenome sequencing was performed at the University of Wisconsin—Madison Biotechnology Center DNA Sequencing Facility (Madison, WI, USA) on the Illumina NovaSeq 6000 (Illumina, Inc., San Diego, CA), resulting in paired-end reads of 2×150 bp. For metagenome sequencing, only two DNA samples from week 8 and Week 18 were selected for sequencing.

### 16S rRNA Amplicon Sequence Analysis

16S rRNA sequencing data was obtained and imported into R version 3.6.1^49^ through RStudio version 1.2.5001, and the packages DADA2^50^ and phyloseq^51^ were used to assign taxonomy and analyze data, respectively. Through the DADA2 package, reads were trimmed using the filterAndTrim function with the following parameters: fruncLen=c(275, 215), maxN=0, truncQ=2, and rm.phix=TRUE. Following read trimming and other quality control, taxonomy was assigned using the assignTaxonomy function of the DADA2 package with the Silva version 132 training data set^52^. For OTUs identified through the assignTaxonomy function that had no domain-level classification given, the Phylum-level classification was manually changed to “Uncharacterized_Bacteria.” All other OTUs that had no Phylum level classification but had a Domain-level classification were manually named to include “Uncharacterized_” in front of the Domain-level classification given for the Phylum-level classification. Unique OTUs for weeks 0, 2, 4 and for weeks 6, 8, 10, 12, 14, 16, 18, 20, 22, 24 were identified if they were observed to occur in 60% or greater of the samples.

### Metagenome Sequencing Data Analysis and Genome Quality Assessment

The two metagenome samples from weeks 8 and 18 were assembled independently. Read trimming was performed with Sickle version 1.33 (https://github.com/najoshi/sickle) using paired-end read trimming with the encoding parameter “-t sanger.” Trimmed reads were assembled with SPAdes v3.12.0^53^ (MetaSPAdes) using the flag --meta and by specifying kmer values of 21, 33, 55, 77, 99, and 127 for assembly. Only scaffolds larger than 1000 bp were considered for downstream analyses. Differential coverage based genome binning was performed using the metawrap^54^ pipeline, making use of two binning softwares: metabat1^55^ and metabat2^56^. To extract and consolidate the best bins produced by each binning software, DASTool^57^ was run on the bins with the flag “--score_threshold 0.4.” Genome completeness and contamination were identified using CheckM^58^ and DASTool. To identify preliminary taxonomy, GTDB-Tk^59^ was run on obtained MAGs using the classify_wf function. The quality of draft genomes was established using MIMAG standards^34^. Open reading frames/Proteins were predicted from the assembled scaffolds using the metagenome mode of Prodigal^60^ (--p meta).

### Concatenated 16 Ribosomal Protein Phylogeny (RP16)

A curated set of publicly available 16 ribosomal protein Hidden Markov Model (HMM) files were utilized to search for ribosomal proteins in the MAGs. All annotated proteins were queried using hmmsearch^61^ with trusted cutoffs for each HMM. A custom python script was used to sort output tables by score and remove duplicate hits to a single genome by retaining hits with higher scores.

To generate a robust concatenated protein alignment of all 16 extracted ribosomal proteins, a manually curated set of these 16 ribosomal proteins for approximately 3500 reference genomes was obtained and imported along with the ribosomal proteins from the MAGs into Geneious Prime. Ribosomal protein sequences were aligned using MAFFT v7.450^62^ (Algorithm: Auto, Scoring matrix: BLOSUM62, Gap open penalty: 1.53, Offset value: 0.123). Geneious Prime was used to concatenate the alignments generated for the 16 ribosomal proteins. All alignment columns that had gaps >= 95% were removed. A preliminary tree was generated with FastTree Version 2.1.11^63^ run with default parameters. This tree was utilized to identify reference genomes that could be removed with little effect on the phylogenetic placement of the MAGs from this study. This resulted in approximately 1500 reference genomes that were used for the final phylogenetic tree. The final tree was implemented in RAxML-HPC v.8^64^ on the CIPRES^65^ platform with the following parameters: bootstopping_type autoMRE, choose_bootstop bootstop, datatype protein, parsimony_seed_val 12345, prot_matrix_spec LG, prot_sub_model PROTCAT. The final tree was visualized in Figtree and iTOL^66^. To generate the phylogenetic tree seen in Figure 6, an unrooted form of the phylogenetic tree was obtained from iTOL. The unedited tree can be accessed at http://itol.embl.de/shared/breister2. The alignment used to generate the phylogenetic tree in phylip format is available in Supplementary Data S1. The unedited phylogenetic tree in newick format is available in Supplementary Data S2.

### Calculation of Replication Rates

All MAGs from the four most abundant phyla *Deltaproteobacteria, Chlorobi, Chloroflexi*, and CPR/*Patescibacteria* were used to estimate replication rates. Trimmed reads were mapped onto genomes using Bowtie2^67^. Read mapping was conducted on each of the genomes, using the corresponding set of reads (e.g. Genomes from week 8 were mapped with reads from week 8) Shrinksam Version 0.9.0 (https://github.com/bcthomas/shrinksam) was used to generate a reduced SAM file containing only reads that mapped to the genome. The sorted SAM files were used as input for iRep^41^, which was run with default parameters for each genome.

### Calculation of Relative Abundance/Coverage of MAGs

Genome Coverage of all MAGs was determined by read mapping using Bowtie2. All mapped SAM files were converted to BAM format using samtools. CoverM (https://github.com/wwood/CoverM) was used to generate read coverage information from the BAM files. An in-house perl script was used to average the depth values for each of the scaffolds within each bin, thereby generating coverage values for each genome bin.

### Determination of Metabolic Potential

The metabolic potential of all MAGs was determined using METABOLIC-C^68^. METABOLIC-C was run on each MAG with reads from weeks 8 or 18 depending on the sample that the MAGs originated from (e.g. MAGs from week 8 were used with reads from week 8). METABOLIC-C was used to generate Supplementary Figures 1 and 2. Resultant metabolic charts produced by METABOLIC-C that involved use of the KEGG database were analyzed by referencing KEGG modules to confirm reactions performed by each step.

### Determining microbial mechanisms of Polymer Composites Breakdown

Several metabolisms proposed to be involved in the degradation of polymer composites are described in Table 1. All metabolic pathways associated with polymer composite degradation were identified using HMM based searches against specific databases (Pfam^69^, KEGG^70^, custom) outlined in Supplementary Table 13. An HMM profile was used to query for the presence of an amidase, AmiE (EC 3.5.1.4), which has been shown to be used in the degradation of polyacrylamide^42^. Also, a set of genes shown to degrade acrylate was identified through MetaCyc^71^, and amino acid fasta files were queried against the genomes using an e-value cutoff of 1e-20.

## Data availability

All DNA sequences (genomes and sequence reads) have been deposited in NCBI Genbank in the BioProject database with accession code PRJNA615962. NCBI Genbank accession numbers for individual genomes are listed in Supplementary Table 4. The authors declare that all other data is available within the article and its supplementary information files, or from the corresponding authors on request.

## Acknowledgements

We would like to acknowledge support from by U.S. Office of Naval Research – Young Investigator Program (ONR-YIP) award [Grant No.: N00014-19-1-2206]. We also thank the University of Wisconsin - Office of the Vice Chancellor for Research and Graduate Education, University of Wisconsin – Department of Bacteriology, and University of Wisconsin – College of Agriculture and Life Sciences for their support. We thank Alejandra Castellanos and Kristopher Kieft, doctoral students from the Prabhakar and Anantharaman labs, respectively, at UW-Madison for setting up preliminary experiments for this study. The authors thank the University of Wisconsin Biotechnology Center DNA Sequencing Facility for providing DNA sequencing facilities and services.

## Contributions

A.B and M.I. carried out the experiments. A.B., Z.Z., and K.A conducted the microbial analyses. M.I., A.B., P.P. and K.A. wrote the manuscript. P.P. and K.A. supervised the project. P.P and K.A. conceived the original idea. All authors discussed the results and contributed to the final manuscript.

## Supplementary Information

**Supplementary Figure 1**. Energy Flow Diagram representing sources of energy at different resolution for the microbial community. Figure was generated with METABOLIC-C.

**Supplementary Figure 2**. Metabolic Connections Diagram representing connectivity in the polymer composite biofilm associated with shared microbial metabolisms. Figure was generated with METABOLIC-C.

**Supplementary Figure 3**. Attenuated total reflection Fourier-transform infrared spectroscopy (ATR-FTIR) graphs of carbon fiber reinforced vinyl ester composites in autoclaved deionized (DI+A) water periodically obtained up to 20 weeks.

**Supplementary Figure 4**. Attenuated total reflection Fourier-transform infrared spectroscopy (ATR-FTIR) graphs of carbon fiber reinforced vinyl ester composites in autoclaved soil solution with deionized (S+DI+A) water periodically obtained up to 20 weeks.

**Supplementary Figure 5**. Estimated growth rates of important microbial groups in the polymer composite biofilm. Each set of boxplots shown describes the growth rate values generated by iRep for genomes originating from the Week 8 metagenome sequencing project (Orange) and Week 18 metagenome sequencing project (Teal). Boxplots were generated using the geom_boxplot() function of the ggplot2 package.

**Supplementary Table 1**. List of OTUs identified in the polymer composite biofilm. The table shows the phylogenetic classification of all OTUS as was generated with DADA2.

**Supplementary Table 2**. Relative Abundance Chart of OTUs identified in the polymer composite biofilm. This chart shows the relative abundances of each of the OTUS within the specified sample. Sample names are listed with “_1” or “_2” at the ends, specifying the duplicate number of the samples.

**Supplementary Table 3**. Abundance and Phylogeny of OTUs in the Polymer Composite Biofilm. This chart describes the OTUs identified by DADA2 within each sample and the relative abundance within that sample of the specific OTU.

**Supplementary Table 4**. Detailed summary of MAG characteristics. The color of the column headers relates to the type of statistic. Classification as a member of CPR/Patescibacteria is given as Y (yes) or N (no). To generate the completion and contamination levels to be used to determine draft genome quality, the set of completion and contamination levels generated by either CheckM or DASTool that resulted in the best draft quality identification was used. If both metrics generated the same draft quality, the set of completion and contamination levels were taken from the program that generated the higher completeness level.

**Supplementary Table 5**. Genome Coverage of all MAGs. This chart shows genome coverage values for all MAGs generated from the respective metagenomes from which they were reconstructed (Week 8 or Week 18).

**Supplementary Table 6**. Summary of presence or absence of individual proteins associated with metabolism. This chart shows the presence (red) or absence (white) of custom HMM profiles from METABOLIC-C ordered by phylum and limited to only MAGs described as high and medium-quality drafts. Bacterial MAGs are grouped by Phylum-level classification and all Archaeal MAGs are grouped together.

**Supplementary Table 7**. Summary of presence or absence of metabolic pathways. This chart generated by METABOLIC-C shows a condensed version of Supplementary Table 5, where the custom HMM profiles from Table S5 are combined into full pathways. Red squares represent presence of a pathway and white squares represent absence of a pathway. Bacterial MAGs are grouped by Phylum-level classification and all Archaeal MAGs are grouped together.Only genomes that were classified as high or medium-quality drafts were used.

**Supplementary Table 8**. Summary of presence or absence of KEGG Modules. This table generated by METABOLIC-C shows the presence (red) or absence (white) of KEGG modules within all MAGs identified as high and medium-quality drafts. Bacterial MAGs are grouped by Phylum-level classification and all Archaeal MAGs are grouped together.

**Supplementary Table 9**. Summary of presence or absence of individual KEGG Module Steps. This table generated by METABOLIC-C shows the presence (red) or absence (white) of KEGG modules, split into pathway steps as identified through KEGG. Bacterial MAGs are grouped by Phylum-level classification and all Archaeal MAGs are grouped together. MAGs are only included if draft quality was identified as medium or high.

**Supplementary Table 10**. FT-IR absorbance (Supplementary Figure 3) obtained from the polymer composite exposed to autoclaved deionized (DI+A) water periodically up to 20 weeks.

**Supplementary Table 11**. FT-IR absorbance (Supplementary Figure 4) obtained from the polymer composite exposed to autoclaved soil solution with deionized water (S + DI + A) periodically up to 20 weeks.

**Supplementary Table 12**. FT-IR absorbance (Figure 7(a)) obtained from the polymer composite exposed to soil solution with deionized water (S + DI) periodically up to 20 weeks.

**Supplementary Table 13**. Details of proteins associated with degradation of polymer composites. This table shows a description of all of the custom protein searches done as shown in Table S4. This chart provides information for accession of the proteins searched along with the source where the proteins were found. Five of the proteins were already queried against the genomes using METABOLIC-C, and those are identified as “Custom HMMs within METABOLIC.” Only genomes that were classified as high or medium-quality drafts were used.

**Supplementary Table 14**. Summary of Presence/Absence of proteins associated with degradation of polymer composites. This chart shows the presence (red) or absence (white) of custom proteins within each of the MAGs described by high or medium draft quality. Bacterial MAGs are grouped by Phylum-level classification and all Archaeal MAGs are grouped together.

**Supplementary Table 15**. Thermogravimetric Analysis (displayed in Figure 3(a)) of Polymer Composite. Sample weight loss with temperature were measured after taken from soil solution with deionized water (S + DI) periodically up to 20 weeks.

**Supplementary Table 16**. Thermogravimetric Analysis (displayed in Figure 3(b)) of Polymer Composite. Sample weight loss with temperature were measured after taken from soil solution with deionized water (S + DI) periodically up to 20 weeks.

**Supplementary Table 17**. Surface nanoindentation data (displayed in Figure 4) obtained from soil solution (S+DI) exposed polymer composite samples periodically up to 24 weeks.

**Supplementary Table 18**. FT-IR Absorbance (shown in Figure 7(b)) obtained from raw FT-IR data (Supplementary Table 10-12) at the wavelength of 1722 cm-1 for C=O bond up to 20 weeks for different environments.

**Supplementary Table 19**. FT-IR Absorbance (presented in Figure 7(c)) collected from raw FT-IR data (Supplementary Table 10-12) at the wavelength of 3400 cm-1 for OH-bond up to 20 weeks for different environments.

**Supplementary Table 20**. Dissolved Organic Carbon content (presented in Figure 8 (b)) measured using the collected solutions from different environmental conditions used in this study and filtered them individually through a 0.45μm pore size filter (Nalgene, 045um syringe filter, 25-mm cellulose acetate membrane).

**Supplementary Data 1**. Concatenated protein alignment used to generate the phylogenetic tree depicted in Figure 6(a).

**Supplementary Data 2**. Newick Tree. The raw text file of the newick format of the phylogenetic tree displayed in Figure 6(a).

## References

1. Krzyzak, A. et al. Sandwich structured composites for aeronautics: methods of manufacturing affecting some mechanical properties. Int. J. Aerosp. Eng. 1–10 (2016).

2. Hota, G. & Liang, R. Advanced Fiber Reinforced Polymer Composites for Sustainable Civil Infrastructures. Int. Symp. Innov. Sustain. od Struct. 2, 234–245 (2011).

3. Allan, M., Thiru, A., Amir, F. & Brahim, B. State-of-the-Art Review on FRP Sandwich Systems for Lightweight Civil Infrastructure. J. Compos. Constr. 21, 4016068 (2017).

4. Girão Coelho, A. M. & Mottram, J. T. A review of the behaviour and analysis of bolted connections and joints in pultruded fibre reinforced polymers. Mater. Des. 74, 86–107 (2015).

5. Keller, T., Schaumann, E. & Vallée, T. Flexural behavior of a hybrid FRP and lightweight concrete sandwich bridge deck. Compos. Part A Appl. Sci. Manuf. 38, 879–889 (2007).

6. Yang, X., Bai, Y. & Ding, F. Structural performance of a large-scale space frame assembled using pultruded GFRP composites. Compos. Struct. 133, 986–996 (2015).

7. Keller, T. Multifunctional and robust composite material structures for sustainable construction Keller, T. Advances in FRP Composites in Civil Engineerin. in Proceedings of the 5th International Conference on FRP Composites in Civil Engineering, CICE 2010 20–25 (2011).

8. Zhu, D. et al. Fiber reinforced composites sandwich panels with web reinforced wood core for building floor applications. Compos. Part B Eng. 150, 196–211 (2018).

9. Cheng, L. & Karbhari, V. M. New bridge systems using FRP composites and concrete: a state-of-the-art review. Prog. Struct. Eng. Mater. 8, 143–154 (2006).

10. Godat, A., Légeron, F., Gagné, V. & Marmion, B. Use of FRP pultruded members for electricity transmission towers. Compos. Struct. 105, 408–421 (2013).

11. Guades, E., Aravinthan, T., Islam, M. & Manalo, A. A review on the driving performance of FRP composite piles. Compos. Struct. 94, 1932–1942 (2012).

12. Lee, S.-B., Rockett, T. J. & Hoffman, R. D. Interactions of water with unsaturated polyester, vinyl ester and acrylic resins. Polymer (Guildf). 33, 3691–3697 (1992).

13. Visco, A. M., Brancato, V. & Campo, N. Degradation effects in polyester and vinyl ester resins induced by accelerated aging in seawater. J. Compos. Mater. 46, 2025–2040 (2011).

14. Wang, J. U. N., Gangarao, H. O. T. A., Liang, R. & Liu, W. Durability and prediction models of fiber-reinforced polymer composites under various environmental conditions: A critical review. J. Reinf. Plast. Compos. 35, 179–211 (2016).

15. Garcia, R., Castellanos, A. & Prabhakar, P. Influence of Arctic seawater exposure on the flexural behavior of woven carbon/vinyl ester composites. J. Sandw. Struct. Mater. 1–19 (2017). doi:10.1177/1099636217710821

16. Singh, A. & Davidson, B. Effects of temperature, seawater and impact on the strength, stiffness, and life of sandwich composites. J. Reinf. Plast. Compos. 30, 269–277 (2011).

17. Scudamore, R. J. & Cantwell, W. J. The effect of moisture and loading rate on the interfacial fracture properties of sandwich structures. Polym. Compos. 23, 406–417 (2002).

18. Siriruk, A., Penumadu, D. & Jack Weitsman, Y. Effect of sea environment on interfacial delamination behavior of polymeric sandwich structures. Compos. Sci. Technol. 69, 821–828 (2009).

19. Loos, A. C., Springer, G. S., Sanders, B. A. & Tung, R. W. Moisture Absorption of Polyester-E Glass Composites. J. Compos. Mater. 14, 142–154 (1980).

20. Pangallo, D. et al. Biodeterioration of epoxy resin: a microbial survey through culture-independent and culture-dependent approaches. Environ. Microbiol. 17, 462–479 (2015).

21. Eliaz, N. et al. Microbial Degradation of Epoxy. Materials (Basel). 11, 2123 (2018).

22. Jecu, L. et al. SUSCEPTIBILITY OF THERMOPLASTIC BASED COMPOSITES TO DEGRADATION BY MICROORGANISMS. Environ. Eng. Manag. J. 14, 2545–2554 (2015).

23. Gu, J.-D., Lu, C., Thorp, K., Crasto, A. & Mitchell, R. Fiber-reinforced polymeric composites are susceptible to microbial degradation. J. Ind. Microbiol. Biotechnol. 18, 364–369 (1997).

24. Wagner, P. A., Little, B. J., Hart, K. R. & Ray, R. I. Biodegradation of composite materials. Int. Biodeterior. Biodegradation 38, 125–132 (1996).

25. Shah, A. A., Hasan, F., Hameed, A. & Ahmed, S. Biological degradation of plastics: A comprehensive review. Biotechnol. Adv. 26, 246–265 (2008).

26. Zyska, B. A short history of investigations of microbial biodeterioration in Poland, 1875– 2002. Int. Biodeterior. Biodegradation 53, 145–150 (2004).

27. Milde, K., Sand, W., Wolff, W. & Bock, E. Thiobacilli of the Corroded Concrete Walls of the Hamburg Sewer System. Microbiology 129, 1327–1333 (1983).

28. Gu, J. D., Ford, T., Thorp, K. & Mitchell, R. Microbial growth on fiber reinforced composite materials. Int. Biodeterior. Biodegrad. 37, 197–204 (1996).

29. Thompson, L. R. et al. A communal catalogue reveals Earth’s multiscale microbial diversity. Nature 551, 457 (2017).

30. Castellanos, A. G., Cinar, K., Guven, I. & Prabhakar, P. Low-Velocity Impact Response of Woven Carbon Composites in Arctic Conditions. J. Dyn. Behav. Mater. (2018). doi:10.1007/s40870-018-0160-8

31. Prabhakar, P. & Garcia, R. Flexural Fatigue Life of Woven Carbon/Vinyl Ester Composites under Sea Water Saturation. arXiv (2019). doi:1911.08665

32. Gewert, B., Plassmann, M. M. & MacLeod, M. Pathways for degradation of plastic polymers floating in the marine environment. Environ. Sci. Process. Impacts 17, 1513–1521 (2015).

33. Nakajima-Kambe, T., Shigeno-Akutsu, Y., Nomura, N., Onuma, F. & Nakahara, T. Microbial degradation of polyurethane, polyester polyurethanes and polyether polyurethanes. Appl. Microbiol. Biotechnol. 51, 134–140 (1999).

34. Bowers, R. M. et al. Minimum information about a single amplified genome (MISAG) and a metagenome-assembled genome (MIMAG) of bacteria and archaea. Nat. Biotechnol. 35, 725–731 (2017).

35. Coates, J. Interpretation of infrared spectra, a practical approach. Encycl. Anal. Chem. Appl. theory Instrum. (2006).

36. Bruce Martin, R. Acid-Catalyzed Ester Hydrolysis. J. Am. Chem. Soc. 89, 2501–2502 (1967).

37. Romera-Castillo, C., Pinto, M., Langer, T. M., Álvarez-Salgado, X. A. & Herndl, G. J. Dissolved organic carbon leaching from plastics stimulates microbial activity in the ocean. Nat. Commun. 9, (2018).

38. Greening, C. et al. Genomic and metagenomic surveys of hydrogenase distribution indicate H2 is a widely utilised energy source for microbial growth and survival. ISME J. 10, 761–777 (2016).

39. Anantharaman, K. et al. Expanded diversity of microbial groups that shape the dissimilatory sulfur cycle. ISME J. 12, 1715–1728 (2018).

40. Morra, M. J. & Dick, W. A. Mechanisms of H2S production from cysteine and cystine by microorganisms isolated from soil by selective enrichment. Appl. Environ. Microbiol. 57, 1413–1417 (1991).

41. Brown, C. T., Olm, M. R., Thomas, B. C. & Banfield, J. F. Measurement of bacterial replication rates in microbial communities. Nat. Biotechnol. 34, 1256–1263 (2016).

42. Kay-Shoemake, J. L., Watwood, M. E., Sojka, R. E. & Lentz, R. D. Polyacrylamide as a substrate for microbial amidase in culture and soil. Soil Biol. Biochem. 30, 1647–1654 (1998).

43. Davey, M. E. & O’toole, G. A. Microbial biofilms: from ecology to molecular genetics. Microbiol. Mol. Biol. Rev. 64, 847–67 (2000).

44. Stipanicev, M. et al. Corrosion behavior of carbon steel in presence of sulfate-reducing bacteria in seawater environment. Electrochim. Acta 113, 390–406 (2013).

45. Hug, L. A. et al. A new view of the tree of life. Nat. Microbiol. 1, 16048 (2016).

46. Anantharaman, K. et al. Thousands of microbial genomes shed light on interconnected biogeochemical processes in an aquifer system. Nat. Commun. 13219, (2016).

47. ASCE’s 2017 Infrastructure Report Card. https://www.infrastructurereportcard.org/ (2020).

48. International., A. ASTM E2546-15 Standard Practice for Instrumented Indentation Testing1. in (ASTM International, 2016).

49. R Core Team. R: A Language and Environment for Statistical Computing. (2019).

50. Callahan, B. J. et al. DADA2: High-resolution sample inference from Illumina amplicon data. Nat. Methods 13, 581–583 (2016).

51. McMurdie, P. J. & Holmes, S. Phyloseq: An R Package for Reproducible Interactive Analysis and Graphics of Microbiome Census Data. PLoS One 8, e61217 (2013).

52. Quast, C. et al. The SILVA ribosomal RNA gene database project: Improved data processing and web-based tools. Nucleic Acids Res. 41, D590–D596 (2013).

53. Nurk, S., Meleshko, D., Korobeynikov, A. & Pevzner, P. metaSPAdes: a new versatile de novo metagenomics assembler. Genome Res. 27, 824–834 (2017).

54. Uritskiy, G. V., DiRuggiero, J. & Taylor, J. MetaWRAP—a flexible pipeline for genome-resolved metagenomic data analysis. Microbiome 6, 158 (2018).

55. Kang, D. D., Froula, J., Egan, R. & Wang, Z. MetaBAT, an efficient tool for accurately reconstructing single genomes from complex microbial communities. PeerJ 3, e1165 (2015).

56. Kang, D. D. et al. MetaBAT 2: An adaptive binning algorithm for robust and efficient genome reconstruction from metagenome assemblies. PeerJ 7, e7359 (2019).

57. Sieber, C. M. K. et al. Recovery of genomes from metagenomes via a dereplication, aggregation and scoring strategy. Nat. Microbiol. 3, 836–843 (2018).

58. Parks, D. H., Imelfort, M., Skennerton, C. T., Hugenholtz, P. & Tyson, G. W. CheckM: Assessing the quality of microbial genomes recovered from isolates, single cells, and metagenomes. Genome Res. 25, 1043–1055 (2015).

59. Chaumeil, P., Hugenholtz, P. & Parks, D. H. GTDB-Tk: a toolkit to classify genomes with the Genome Taxonomy Database. Bioinformatics btz848, 1–3 (2019).

60. Hyatt, D. et al. Prodigal: Prokaryotic gene recognition and translation initiation site identification. BMC Bioinformatics 11, (2010).

61. Eddy, S. R. Accelerated profile HMM searches. PLoS Comput. Biol. 7, e1002195 (2011).

62. Katoh, K. & Standley, D. M. MAFFT multiple sequence alignment software version 7: Improvements in performance and usability. Mol. Biol. Evol. 30, 772–780 (2013).

63. Price, M. N., Dehal, P. S. & Arkin, A. P. Fasttree: Computing large minimum evolution trees with profiles instead of a distance matrix. Mol. Biol. Evol. 26, 1641–1650 (2009).

64. Stamatakis, A. RAxML version 8: A tool for phylogenetic analysis and post-analysis of large phylogenies. Bioinformatics 30, 1312–1313 (2014).

65. Miller, M. A., Pfeiffer, W. & Schwartz, T. Creating the CIPRES Science Gateway for inference of large phylogenetic trees. in 2010 Gateway Computing Environments Workshop (GCE) 1–8 (2010). doi:10.1109/GCE.2010.5676129

66. Letunic, I. & Bork, P. Interactive Tree Of Life (iTOL) v4: recent updates and new developments. Nucleic Acids Res. 47, W256–W259 (2019).

67. Langmead, B. & Salzberg, S. L. Fast gapped-read alignment with Bowtie 2. Nat. Methods 9, 357–359 (2012).

68. Zhou, Z., Tran, P., Liu, Y., Kieft, K. & Anantharaman, K. METABOLIC: A scalable high-throughput metabolic and biogeochemical functional trait profiler based on microbial genomes. bioRxiv 761643 (2019). doi:10.1101/761643

69. El-Gebali, S. et al. The Pfam protein families database in 2019. Nucleic Acids Res. 47, D427–D432 (2019).

70. Kanehisa, M. & Susuma, G. KEGG: Kyoto Encyclopedia of Genes and Genomes. Nucleic Acids Res. 28, 27–30 (2000).

71. Caspi, R. et al. The MetaCyc database of metabolic pathways and enzymes. Nucleic Acids Res. 46, D633–D639 (2018).

72. Scott, T. F., Cook, W. D. & Forsythe, J. S. Kinetics and network structure of thermally cured vinyl ester resins. Eur. Polym. J. 38, 705–716 (2002).

