## Supplementary Figure 1 for "Microbial dark matter driven degradation of carbon fiber polymer composites"

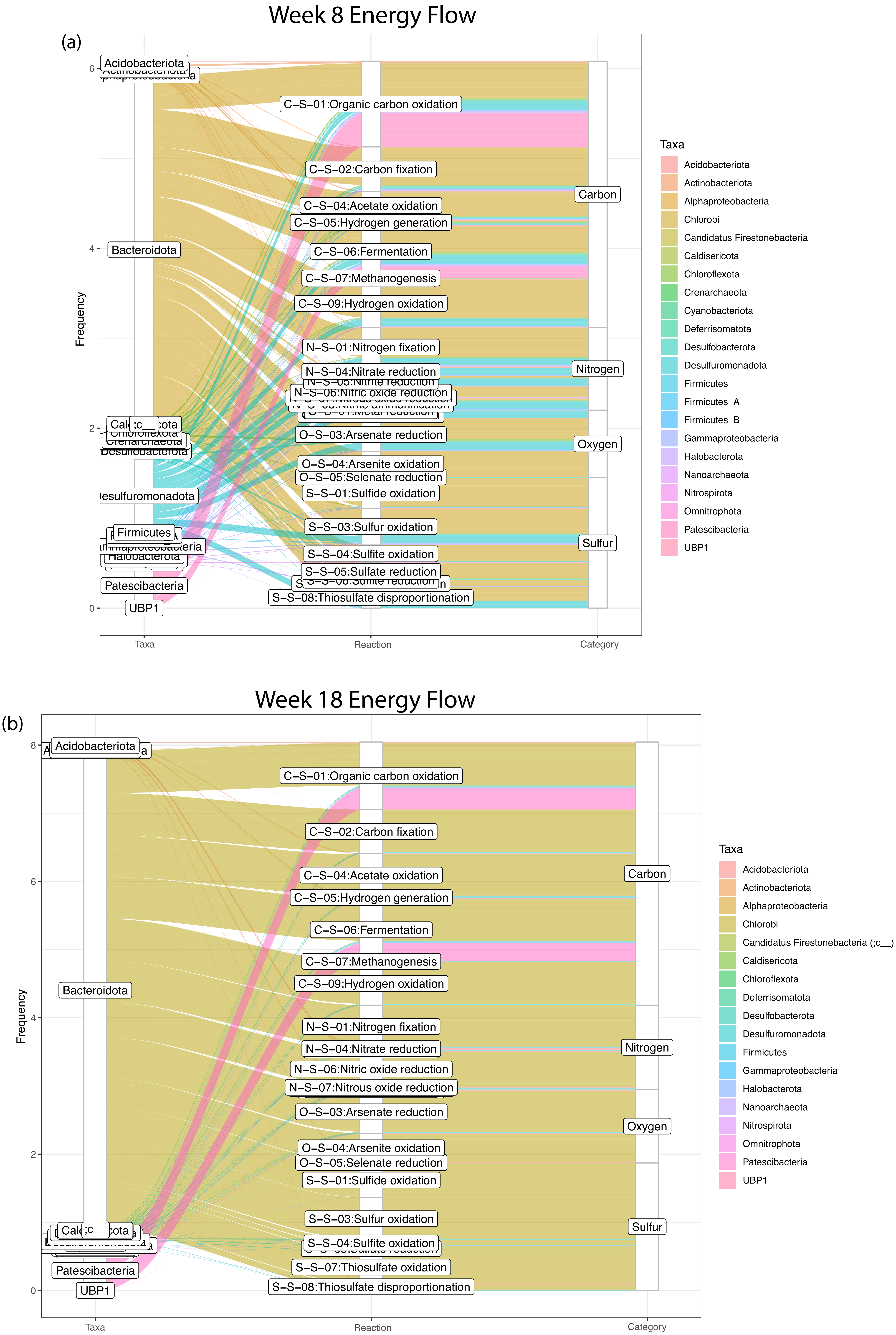


**Supplementary Figure 1.** The charts shown in this figure show the METABOLIC-C-generated energy flow diagrams for genomes from the week 8 sample (a) and the week 18 (a) sample. This chart relates phylogeny and the reactions performed by those organisms to their relative abundance within four metabolic categories. To generate this plot, genomes from week 8 had the week 8 sequencing reads mapped to them and the genomes from week 18 had the week 18 sequencing reads mapped to them.
