## Supplementary Figure 2 for "Microbial dark matter driven degradation of carbon fiber polymer composites"

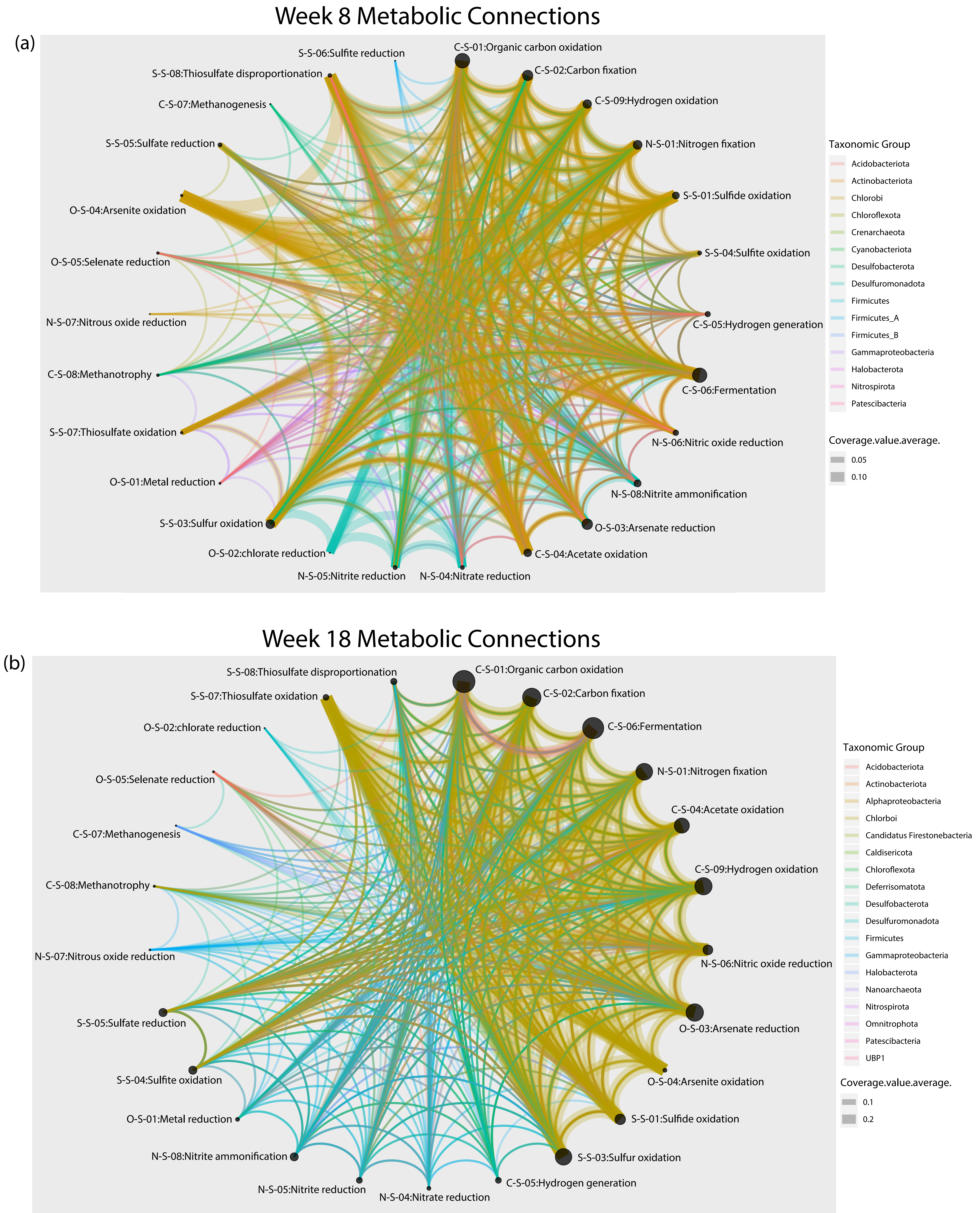


**Supplementary Figure 2.** This chart shows the metabolic connections between metabolic categories generated for metagenomes from week 8 of incubation (a) and week 18 of incubation (b). As with the metabolic energy flow diagrams shown, the charts were generated with METABOLIC-C using read mapping. Genomes from a specific sample (Week 8 or Week 18) had the corresponding set of sequencing reads mapped to them to generate these charts.
