## Supplementary figures and images for "Microbial dark matter driven degradation of carbon fiber polymer composites"

### Supplementary Figure 3

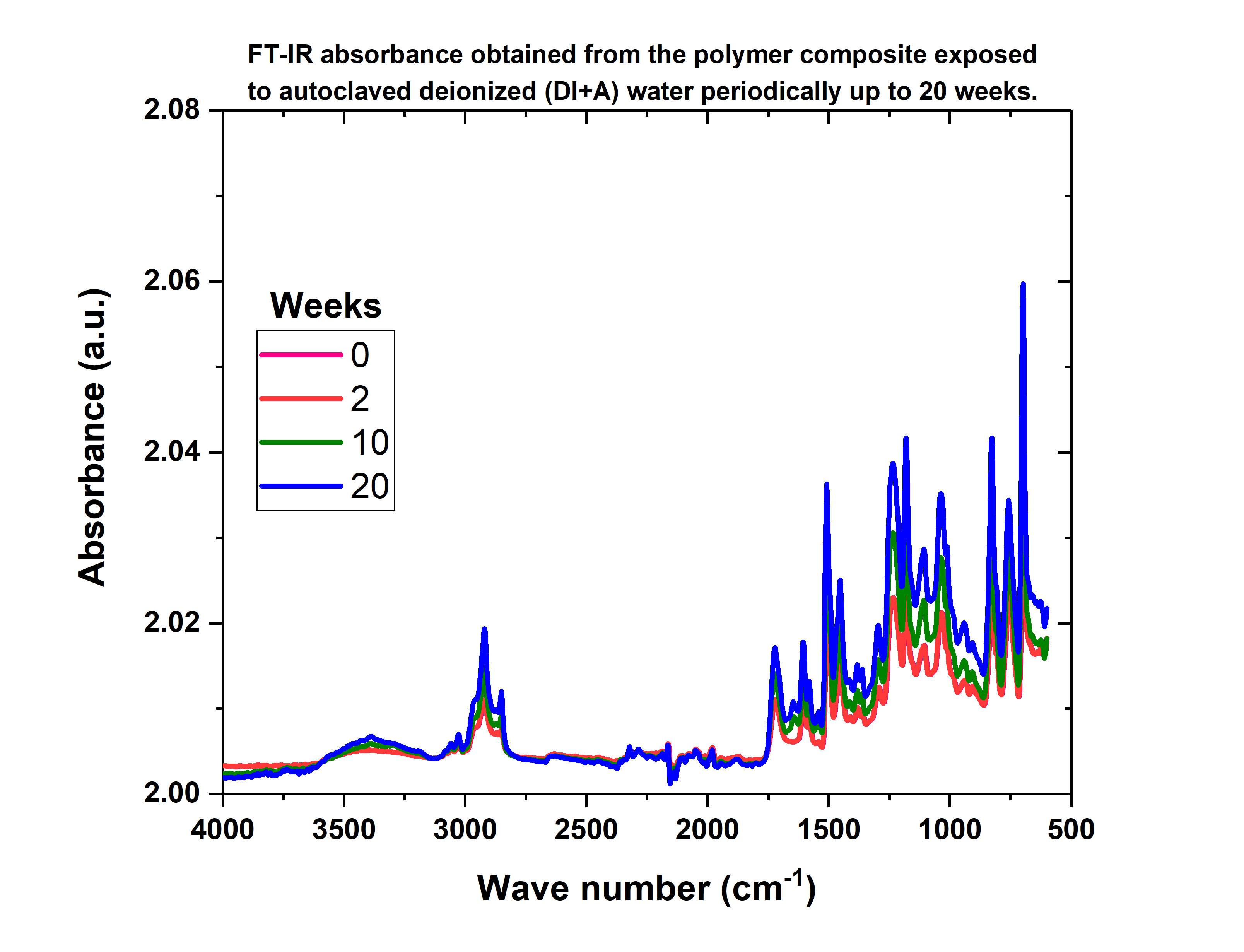

### Supplementary Figure 4

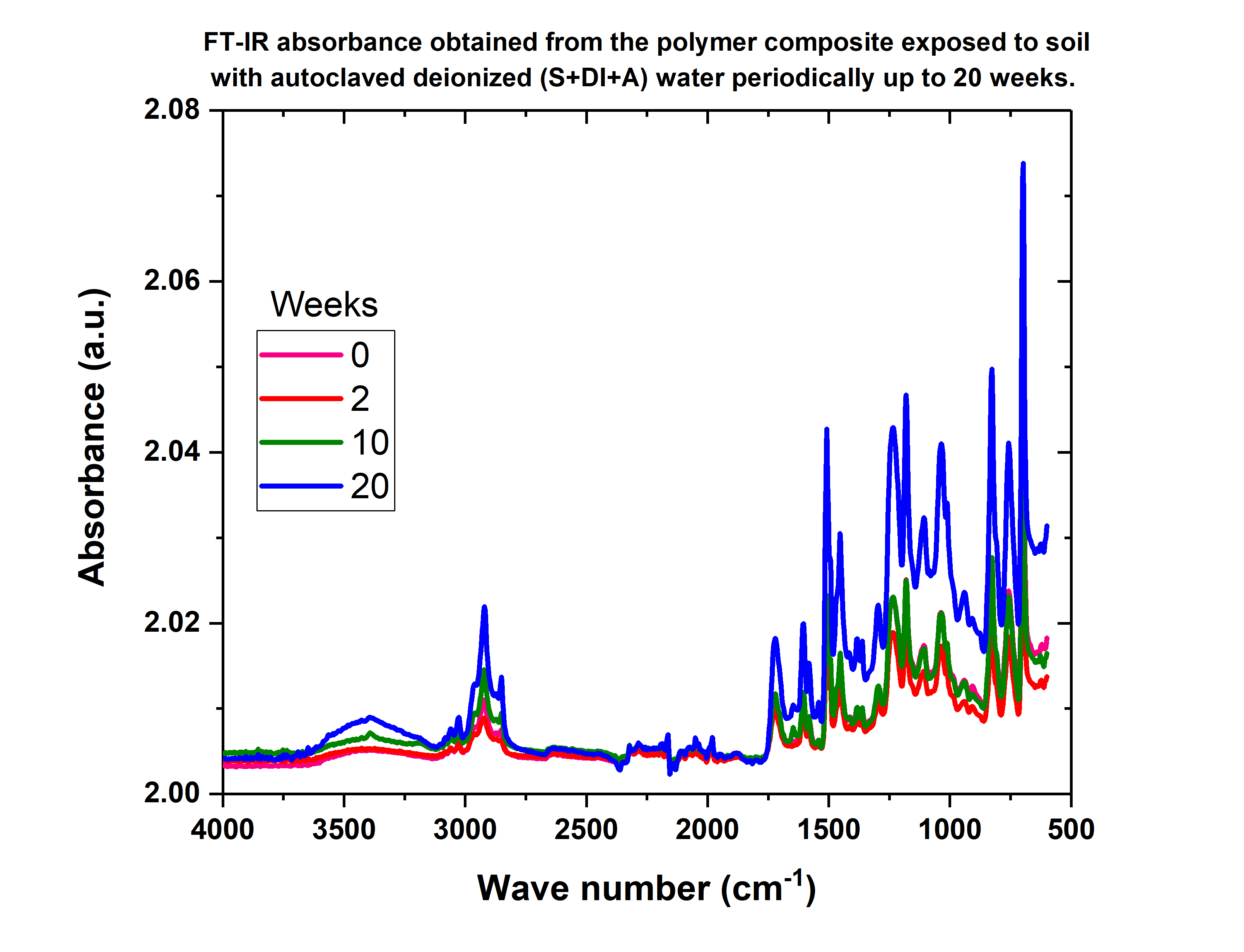
