## Supplementary Figure 5 for "Microbial dark matter driven degradation of carbon fiber polymer composites"

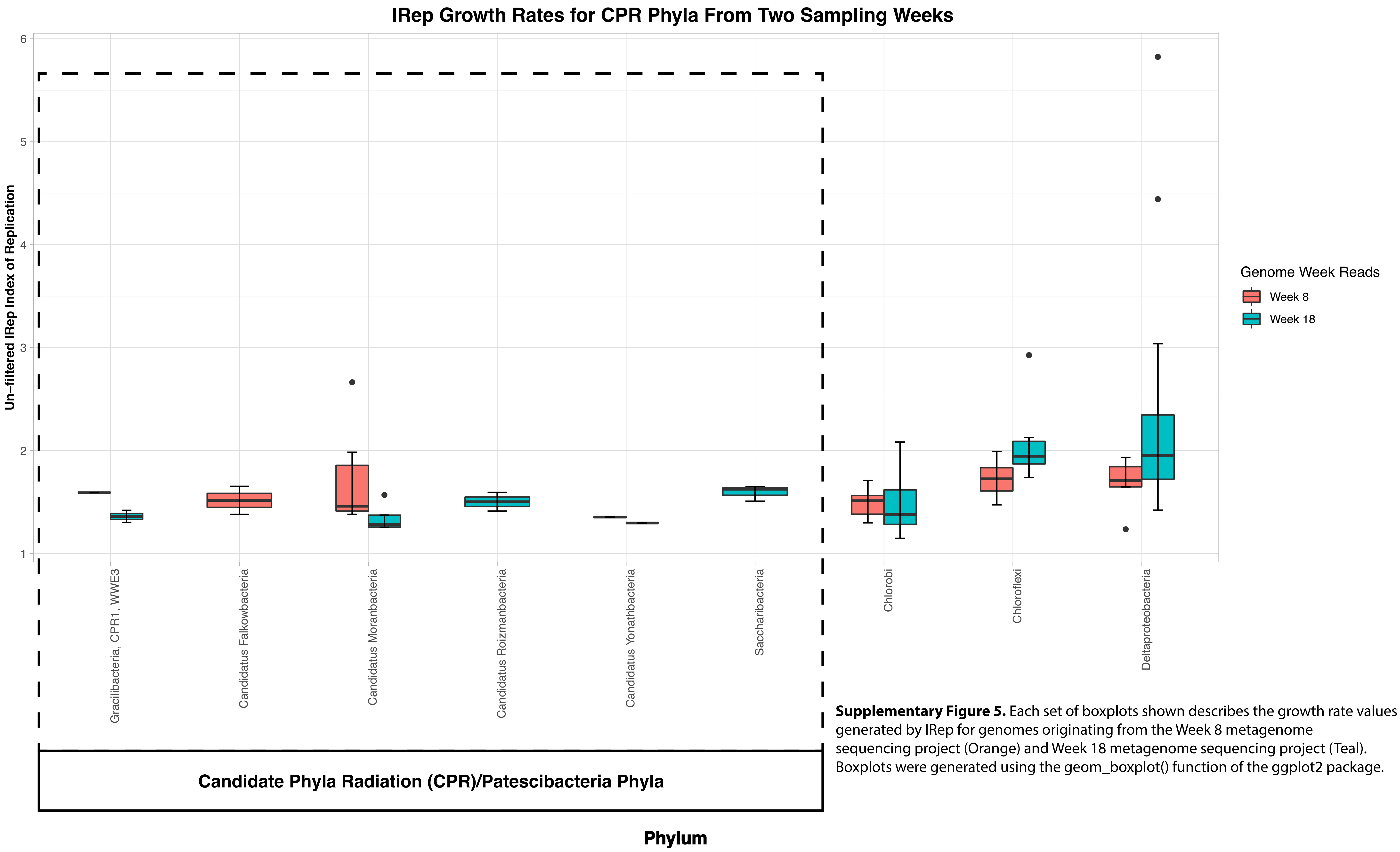


**Supplementary Figure 5.** Each set of boxplots shown describes the growth rate values generated by IRep for genomes originating from the Week 8 metagenome sequencing project (Orange) and Week 18 metagenome sequencing project (Teal). Boxplots were generated using the geom_boxplot() function of the ggplot2 package.
